# A deterministic, c-di-GMP-dependent genetic program ensures the generation of phenotypically similar, symmetric daughter cells during cytokinesis

**DOI:** 10.1101/2024.02.06.579105

**Authors:** María Pérez-Burgos, Marco Herfurth, Andreas Kaczmarczyk, Andrea Harms, Katrin Huber, Urs Jenal, Timo Glatter, Lotte Søgaard-Andersen

**Affiliations:** Department of Ecophysiology, Max Planck Institute for Terrestrial Microbiology, Marburg, Germany; Biozentrum, University of Basel, Basel, Switzerland; Core Facility for Mass Spectrometry & Proteomics, Max Planck Institute for Terrestrial Microbiology, Marburg, Germany

**Keywords:** c-di-GMP, diguanylate cyclase, phosphodiesterase, divisome, polarity, type IV pili, type IV pili-dependent motility, gliding motility, heterogeneity, cell cycle regulation

## Abstract

Phenotypic heterogeneity in bacteria results from stochastic processes or deterministic genetic programs. These deterministic programs often incorporate the versatile second messenger c-di-GMP, and by deploying c-di-GMP metabolizing enzyme(s) asymmetrically during cell division give rise to daughter cells with different c-di-GMP levels. By contrast, less is known about how phenotypic heterogeneity is kept to a minimum. Here, we identify a deterministic c-di-GMP-dependent genetic program that is hardwired into the cell cycle of *Myxococcus xanthus* to minimize phenotypic heterogeneity and guarantee the formation of phenotypically similar daughter cells during division. Cells lacking the diguanylate cyclase DmxA have an aberrant motility behaviour. DmxA is recruited to the cell division site and its activity switched on during cytokinesis, resulting in a dramatic but transient increase in the c-di-GMP concentration. During cytokinesis, this c-di-GMP burst ensures the symmetric incorporation and allocation of structural motility proteins and motility regulators at the new cell poles of the two daughters, thereby generating mirror-symmetric, phenotypically similar daughters with correct motility behaviours. These findings suggest a general c-di-GMP-dependent mechanism for minimizing phenotypic heterogeneity, and demonstrate that bacteria by deploying c-di-GMP metabolizing enzymes to distinct subcellular locations ensure the formation of dissimilar or similar daughter cells.

## Introduction

In bacteria, motility and its regulation contribute to colonization of hosts and other habitats, biofilm formation, virulence and predation ^1^. The ubiquitous second messenger cyclic di-GMP (c-di-GMP) is a key regulator of bacterial motility ^2–4^. Generally, high c-di-GMP levels inhibit flagella-based swimming motility and stimulate surface adhesion and type IV pili (T4P)-dependent surface motility, thereby promoting surface colonization, biofilm formation and virulence ^2–4^. Several bacterial species that alternate between planktonic and surface-adhered lifestyles harness this duality of c-di-GMP – inhibition of flagella-based motility and stimulation of surface adhesion and T4P-based motility – to establish deterministic genetic programs that are hardwired into the cell cycle to produce phenotypically distinct daughter cells during cell division, thereby generating phenotypic heterogeneity within a population of genetically identical cells ^4, 5^. Here, we report that a c-di-GMP-based, deterministic genetic program is hardwired into the cell cycle of *Myxococcus xanthus* to minimize phenotypic heterogeneity and guarantee the formation of phenotypically similar daughter cells during division.

*M. xanthus* does not have flagella and only translocates on surfaces. For this, the rod-shaped cells use two motility systems, one for T4P-dependent motility and one for gliding ^6^. Motility and its regulation are important for the social behaviours of *M. xanthus,* including the formation of spreading colonies in which cells prey on other microbes and spore-filled fruiting bodies in the absence of nutrients ^7^. Both motility systems are highly polarized, i.e. the core T4P machine (T4PM) is present at both cell poles ^8–11^ but only active at one pole at a time ensuring unipolar T4P assembly ^12^, and the Agl/Glt machine for gliding only assembles at one pole at a time ^13–18^. Because the two machines are active at the same pole, cells translocate unidirectionally across surfaces with a piliated leading and non-piliated lagging cell pole ^19–21^. To regulate their social behaviours, *M. xanthus* cells occasionally reverse their direction of movement ^22^. The Frz chemosensory system induces reversals ^22^, which at the cellular level entails the inversion of the polarity of the two motility machines ^6^.

The so-called polarity module establishes the unipolar assembly of the active motility machines. The output of this module is generated by the small GTPase MglA, a nucleotide-dependent molecular switch that is inactive in its GDP-bound and active in its GTP-bound state ^23, 24^. GTP-bound MglA localizes to the leading pole ^23, 24^ where it interacts with effectors to stimulate T4P formation and assembly of the Agl/Glt machine ^18, 20, 21, 25^. The bipartite RomR/RomX and MglB/RomY complexes of the polarity module regulate the nucleotide-bound state and bring MglA-GTP to the leading pole by functioning as a guanine nucleotide exchange factor (GEF) ^26^ and a GTPase activating protein (GAP) ^23, 24, 27^, respectively. These four proteins and the MglC adaptor protein also localize asymmetrically to the poles ^23, 24, 26–29^. During reversals, the Frz system acts on the polarity module to induce an inversion of its polarity ^30, 31^, laying the foundation for activating the motility machineries at the new leading pole. The asymmetric polar localization of the polarity proteins relies on an intricate set of positive and negative feedback loops between the six proteins ^23, 24, 29, 32^. Consequently, mutants lacking one or more polarity proteins have an abnormal localization of the remaining proteins, and such mutants are either non-motile or have an abnormal motility behaviour, i.e. hyper-reverse independently of the Frz system or hypo-reverse and are non-responsive to Frz signalling ^18, 23, 24, 26, 27, 29, 33^.

The c-di-GMP level is determined by the opposing activities of diguanylate cyclases (DGCs), which contain the catalytic GGDEF domain and synthesize c-di-GMP, and phosphodiesterases (PDEs), which degrade c-di-GMP ^2, 3^. *M. xanthus* encodes 11 GGDEF domain proteins predicted to have DGC activity ^34^. The systematic inactivation of the 11 corresponding genes identified DmxA as the only DGC implicated in motility during growth ^34^, while the DGC DmxB is specifically important for fruiting body formation ^35^. The functions of the remaining nine DGCs are not known.

Here, we addressed the mechanism of DmxA. We report that DmxA is recruited to the cell division site, and its DGC activity switched on late during cytokinesis, resulting in a dramatic but transient increase in the c-di-GMP concentration. The burst in c-di-GMP ensures the equal incorporation and allocation of structural motility proteins and polarity proteins at the new cell poles of the two daughters, thereby generating mirror-symmetric, phenotypically similar daughters with correct motility behaviour. Thus, for the first time, evidence is provided that, during cell division, c-di-GMP guarantees the generation of phenotypically similar offspring.

## Results

### DmxA is a dimeric DGC with a low-affinity I-site

Based on sequence analysis, DmxA has an N-terminal transmembrane domain (TMD) with six α-helices, followed by two GAF domains, and the catalytic GGDEF domain with the active (A)-site and a c-di-GMP-binding inhibitory (I)-site (Fig. 1a), which is involved in allosteric feedback inhibition of activity in other DGCs ^36^. A His_6_-DmxA variant comprising the two GAF domains and the GGDEF domain is enzymatically active and binds c-di-GMP *in vitro* ^34^. To understand how the different domains contribute to catalytic activity, we purified five soluble MalE-tagged DmxA variants (Fig. 1b; S1a-b), i.e. variants containing the two GAF domains and the GGDEF domain (MalE-DmxA^WT^), only the two GAF domains (MalE-DmxA^GAF×2^), only the GGDEF domain (MalE-DmxA^GGDEF^), and MalE-DmxA^WT^ variants with substitutions in either the catalytic site (MalE-DmxA^E626A^) or the I-site (MalE-DmxA^R615A^). In size exclusion chromatography (SEC), all variants except MalE-DmxA^GGDEF^ eluted at sizes corresponding to dimers, while MalE-DmxA^GGDEF^ eluted at a size corresponding to a monomer (Fig. 1b; S1a), indicating that the GAF domain-containing region is important for dimer formation. Indeed, a high-confidence Alphafold-Multimer structural model supports that DmxA forms a symmetric dimer in which the protomers interact extensively *via* two α-helices connecting GAF1 to GAF2, and GAF2 to the GGDEF domain and in which the two A-sites are in close proximity and facing each other (Fig. 1c; S1c). Consistently, active DGCs are dimeric, dimerization is mediated by domain(s) outside of the GGDEF domain ^37, 38^, and in solved structures of DGCs the two A-sites are in close proximity ^37–40^.

**Figure 1.**
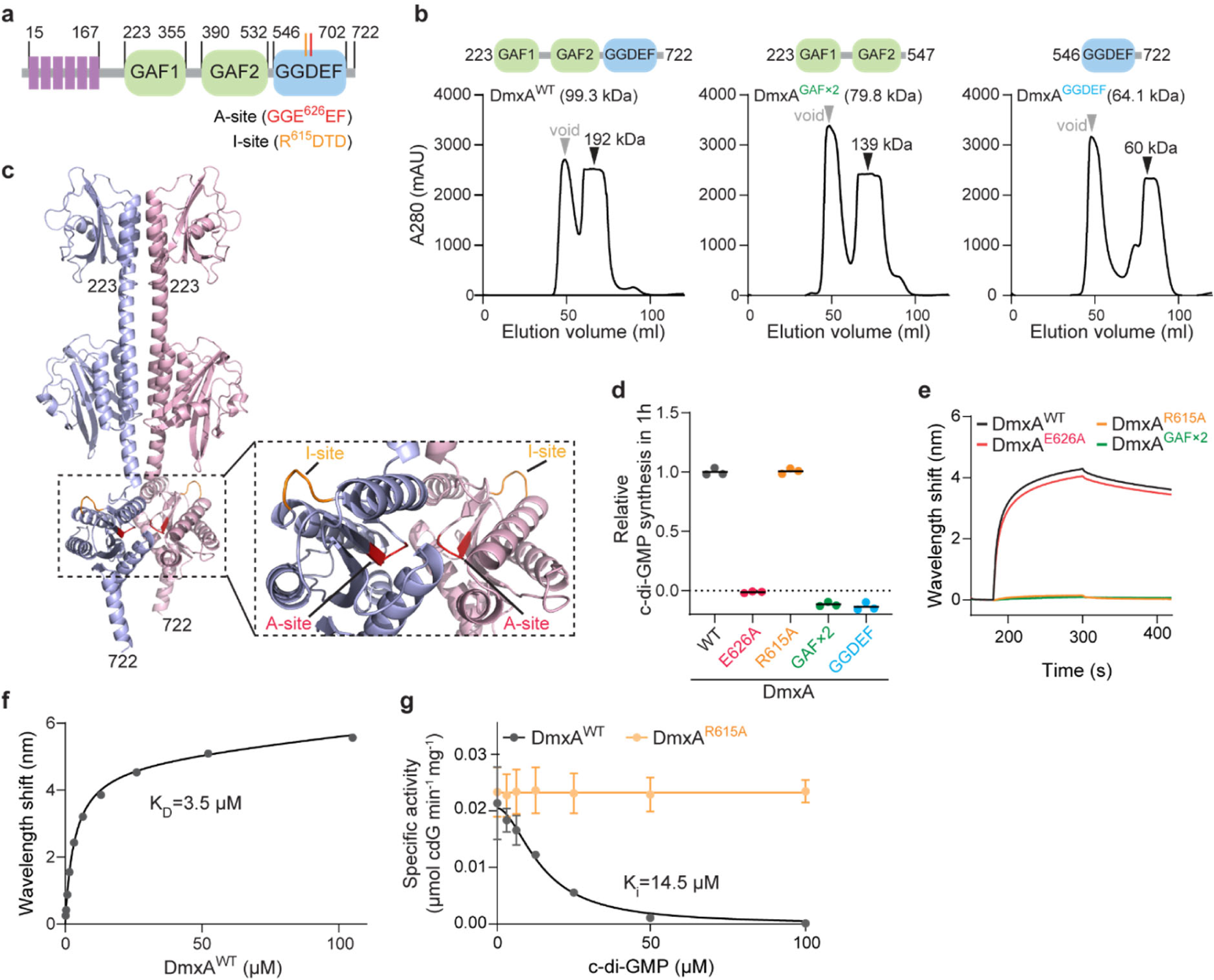
DmxA has DGC activity and a low-affinity I-site. **a**, Domain architecture of DmxA. **b**, SEC of MalE-DmxA variants. Domain architecture of DmxA variants are shown above chromatograms. Grey arrowheads indicate void volume, and black arrowheads elution volume with the corresponding calculated molecular weight. **c**, AlphaFold-Multimer structural model of dimeric DmxA. The transmembrane helices were removed before generating the model, residue numbers are indicated. **d**, *In vitro* DGC assay of the indicated MalE-DmxA variants. The relative amount of c-di-GMP synthesized after 1 h was determined by measuring released inorganic pyrophosphate. Measurements from three technical replicates are shown relative to the mean (black lines) of MalE-DmxA^WT^. **e**, Bio-Layer Interferometric analysis of c-di-GMP binding by MalE-DmxA variants. Streptavidin-coated sensors were loaded with 500 nM biotinylated c-di-GMP and probed with 10 µM of the indicated proteins. The interaction kinetics were followed by monitoring the wavelength shift during the association and dissociation of the analyte. **f**, Determination of K_D_ of MalE-DmxA for c-di-GMP. Plot shows the equilibrium levels measured at the indicated MalE-DmxA^WT^ concentrations (see also Fig S1d). The data were fitted to a non-cooperative one-site specific-binding model. **g**, Determination of K_i_ of MalE-DmxA for c-di-GMP. Inhibition of the specific activity of DmxA^WT^ and DmxA^R615A^DGC activity over time was measured as in **d** in the presence of different c-di-GMP concentrations. Points and error bars represent the mean ± standard deviation (SD) calculated from three biological replicates. The data were fitted to an inhibition model with variable slope.

MalE-DmxA^WT^ and MalE-DmxA^R615A^ had DGC activity, while MalE-DmxA^E626A^ and MalE-DmxA^GAF×2^, as expected, did not (Fig. 1d). Monomeric MalE-DmxA^GGDEF^ did not have DGC activity (Fig. 1d), providing additional support that the GAF domain-containing region is important for dimerization and, therefore, DGC activity. In Bio-Layer Interferometry, MalE-DmxA^WT^ and MalE-DmxA^E626A^ bound c-di-GMP, while MalE-DmxA^R615A^, as expected, and MalE-DmxA^GAF×2^ did not; we determined a K_D_ of 3.5 µM for MalE-DmxA^WT^ (Fig. 1e-f; S1d). C-di-GMP did not significantly inhibit MalE-DmxA^R615A^, while MalE-DmxA^WT^ was inhibited in a cooperative manner (n_h_ = 1.8), with an inhibitory constant (K_i_) of ∼15 µM (Fig. 1g). This concentration is not only ∼10-fold higher than the c-di-GMP concentration (1.4±0.5 µM) measured in an unsynchronized population of *M. xanthus* cells ^41^ but also significantly higher than in other DGCs, e.g. DgcA and PleD of *C. crescentus* have K_i_‘s of 1 µM and 6 µM, respectively ^36^.

We conclude that DmxA has DGC activity and a low-affinity I-site. Moreover, the GAF domain-containing region is important for dimerization, as described for eukaryotic PDEs with an analogous domain architecture ^42^.

### The Δ*dmxA* mutant has an aberrant motility behaviour with aberrant reversals and a cell polarity defect

A disruption of *dmxA* by a plasmid insertion (Ω*dmxA*) caused reduced T4P-dependent motility ^34^. We generated an in-frame deletion mutation of *dmxA* (Δ*dmxA*) to understand the underlying mechanism. In population-based motility assays on 0.5% agar, which is most favourable for T4P-dependent motility, wild-type (WT) generated expanding colonies with the characteristic flares, while the Δ*pilA* negative control, which lacks the major pilin of T4P ^43^, did not (Fig. 2a). The Δ*dmxA* mutant displayed significantly reduced colony expansion, which was restored upon complementation (Fig. 2a). A Δ*dmxA*Δ*gltB* double mutant, which lacks a component of the Agl/Glt machine ^13^, also had significantly reduced colony expansion compared to the Δ*gltB* mutant (Fig. 2a). Thus, Δ*dmxA* causes a T4P-dependent motility defect. On 1.5% agar, which is most favourable for gliding, WT displayed the characteristic single cells at the colony edge, while the Δ*gltB* negative control did not (Fig. 2a). The Δ*dmxA* mutant had single cells at the colony edge but significantly reduced colony expansion, which was restored upon complementation (Fig. 2a). A Δ*dmxA*Δ*pilA* double mutant had single cells at the colony edge but a more significant expansion defect than the Δ*pilA* mutant (Fig. 2a). Thus, the Δ*dmxA* mutation also causes a gliding defect. We previously reported that the Ω*dmxA* mutant only had a defect in T4P-dependent motility ^34^. However, in those experiments, we exclusively focused on the presence or absence of single cells at the colony edge and not overall colony expansion.

**Figure 2.**
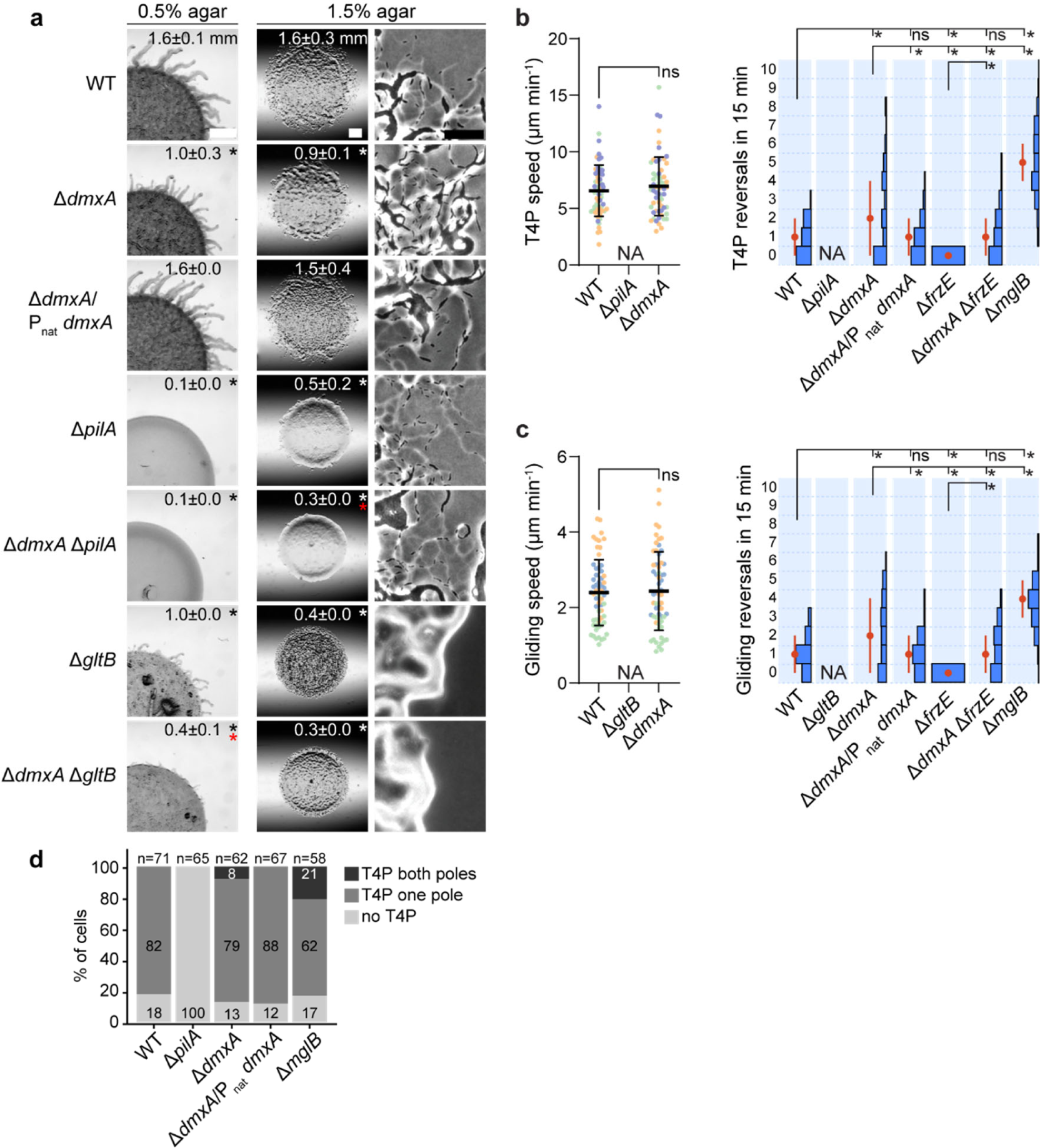
The Δ*dmxA* mutant has an aberrant motility behaviour with aberrant reversals and a cell polarity defect. **a**, Population based motility assays. T4P-dependent motility and gliding were analysed on 0.5% and 1.5% agar, respectively. Numbers indicate the colony expansion in 24 h as mean ± SD (n=3 biological replicates). In the complementation strain, *dmxA* was expressed from its native promoter from a plasmid integrated in a single copy at the Mx8 *attB* site. Black, white and red asterisks indicate *p*<0.05, Student’s *t*-test against WT (black, white) or the Δ*pilA* or Δ*gltB* controls (red). Scale bars, 1 mm (left), 1 mm (middle), 50 μm (right). **b-c**, Single cell-based motility assays. T4P-dependent motility was measured for cells on a polystyrene surface covered with 1% methylcellulose (b) and gliding on 1.5% agar supplemented with 0.5% CTT (c). Cells were imaged for 15 min with 30 s intervals. Speed (n=20 in each of three biological replicates indicated in different colours) and number of reversals (n=30 in each of three biological replicates). Only cells moving during the entire recording interval were included. For speed, error bars represent mean ± SD calculated from all data points. Reversals are represented as histograms based on all three replicates, error bars indicate the median ± median absolute deviation (MAD). NA, not applicable because cells are non-motile. **\*** *p*<0.05, ns, not significant; statistical tests: speed, Mann-Whitney test, reversals, one-way ANOVA multiple comparison test and Fishers LSD test. **d**, Quantification of T4P localization based on transmission electron microscopy. Total number of cells from at least three biological replicates indicated above.

We assayed single-cell motility to uncover how the Δ*dmxA* mutation causes motility defects (Fig. 2b-c). Δ*dmxA* cells had the same speed as WT but reversed aberrantly and had a much broader distribution of reversal frequencies than WT, ranging from cells that did not reverse to cells that reversed up to eight times during the experiment. The reversal defect was corrected upon complementation. Moreover, while Δ*frzE* cells, which lack the FrzE kinase essential for Frz-induced reversals ^44^, did not reverse, many Δ*dmxA*Δ*frzE* cells still reversed. Mutants with Frz-independent reversals generally lack polar MglB/RomY GAP activity, causing MglA-GTP to localize to both cell poles ^18, 23, 24, 27^. Interestingly, the Δ*dmxA* mutant also had a much broader distribution of reversal frequencies than the Δ*mglB* mutant, which lacks MglB/RomY GAP activity, causing essentially all cells to reverse or hyper-reverse.

Because lack of GAP activity results in T4P formation at both cell poles in many cells ^21^, we determined the piliation pattern of Δ*dmxA* cells using transmission electron microscopy. 82% of WT cells were unipolarly piliated and the remaining cells unpiliated, while 21% of Δ*mglB* cells were bipolarly and 62% unipolarly piliated (Fig. 2d). Importantly, 8% of Δ*dmxA* cells were bipolarly and 79% unipolarly piliated (Fig. 2d; S2a).

The structural proteins of the two motility machines, the Frz proteins, and the polarity proteins accumulated at the same level in WT and the Δ*dmxA* mutant (Fig. S2b-g), supporting that changes in protein accumulation are not responsible for the aberrant motility behaviour.

We conclude that cells lacking DmxA have motility defects caused by aberrant reversals and have an unusually broad distribution of reversal frequencies. Moreover, the Δ*dmxA* mutant has a cell polarity defect, and the underlying mechanism differs from that of a mutant lacking MglB/RomY GAP activity.

### DmxA is recruited to the division site by the divisome late during cytokinesis

To understand how DmxA impacts motility behaviour and cell polarity, we determined the localization of an active (see below) DmxA-mVenus fusion expressed from the native site. Remarkably, DmxA-mVenus localized at mid-cell in ∼5% of cells, while the remaining cells had a speckle-like pattern along the cell body (Fig. 3a). In all cells with a mid-cell cluster, the cluster co-localized with a cell division constriction. Still, not all constricting cells had a DmxA-mVenus cluster, suggesting that DmxA-mVenus is recruited to the division site late during cytokinesis. Indeed, we observed by time-lapse microscopy (Fig. 3b) that (i) DmxA-mVenus cluster formation at the division site was visible in each division event, (ii) the cluster appeared at the division site 20±15 min before completion of cytokinesis, and (iii) disintegrated upon completion of cytokinesis. Consistently, the mean cluster lifetime was 20±16 min (Fig. 3b). The 5 h doubling time of *M. xanthus* and this cluster lifetime agree well with the percentage of cells with a mid-cell cluster in an unsynchronized cell population.

**Figure 3.**
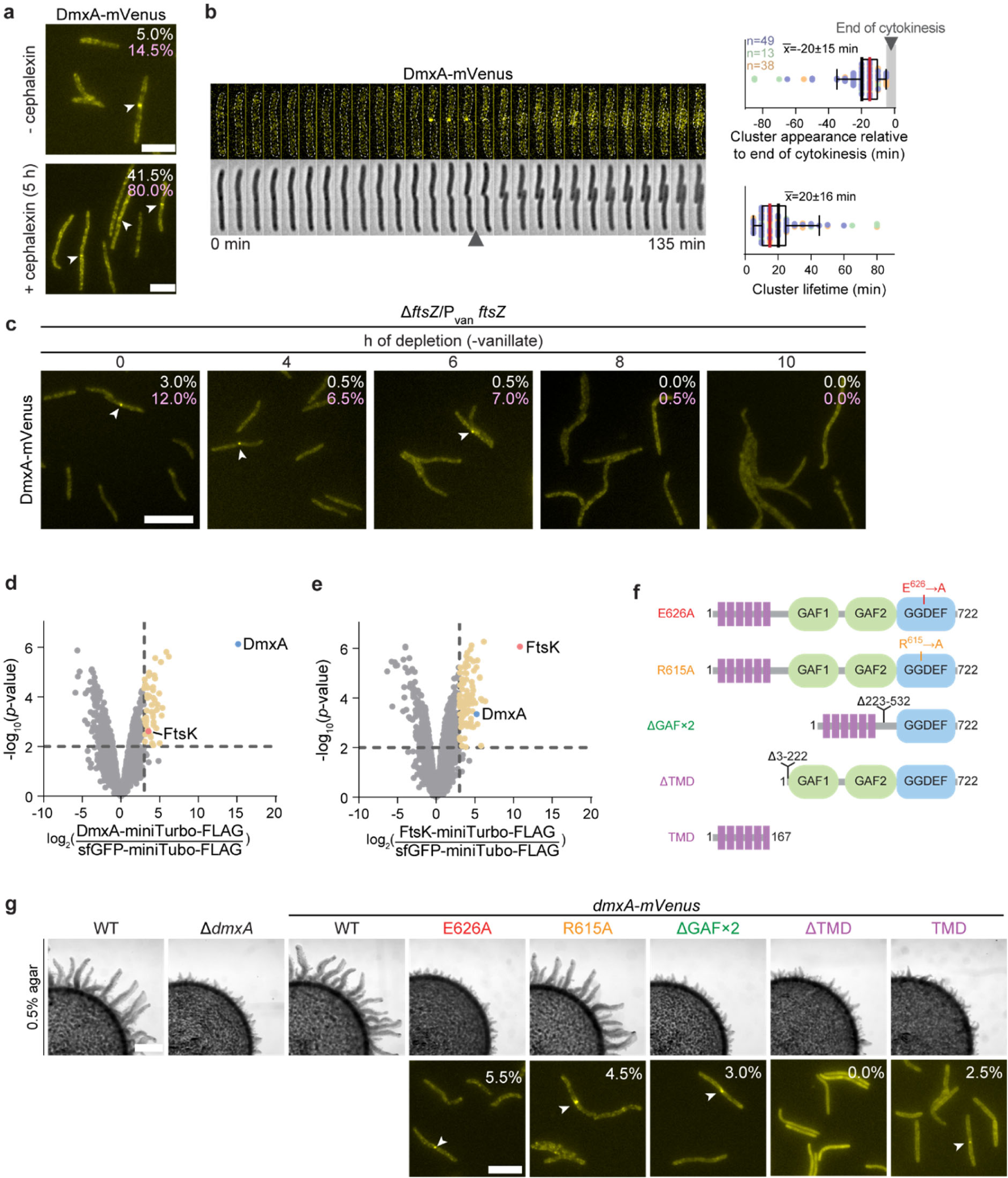
DmxA is recruited to the division site during cytokinesis by the divisome and its function depends on DGC activity. **a**, Localization of DmxA-mVenus in the presence and absence of cephalexin. The percentage of cells with a DmxA-mVenus cluster (white) or a constriction (pink) is indicated (n=200 from one biological replicate). Scale bars, 5µm. **b**, DmxA-mVenus localization during the cell cycle. Left panels, epifluorescence and phase-contrast images from time-lapse microscopy of cells expressing DmxA-mVenus. Images were recorded every 5 min; black arrowhead indicates completion of cytokinesis (defined as the first frame at which daughters were clearly separated); right panels, analysis of the appearance relative to completion of cytokinesis and lifetime of DmxA-mVenus clusters. The first time point after completion of cytokinesis is defined as t=0 and indicated by the grey vertical bar. Box plots show the median (red) and mean (black) with upper and lower quartiles and whiskers present 10^th^ and 90^th^ percentile; n=100 from three biological replicates; number of cells per replicate and the corresponding data points are in matching colours. **c**, Localization of DmxA-mVenus during FtsZ depletion. Cells were grown in the presence of 10 µM vanillate before starting the depletion. The percentage of cells with a cluster (white) or a constriction (pink) are indicated (n=200 from one biological replicate). White arrowheads indicate DmxA-mVenus clusters. Scale bar, 10 µm. **d-e**, Proximity labelling using DmxA-miniTurbo-FLAG or FtsK-miniTurbo-FLAG as baits compared to sfGFP-miniTurbo-FLAG. Volcano plots show proteins enriched by DmxA-miniTurbo-FLAG (d) and FtsK-miniTurbo-FLAG (e). DmxA-miniTurbo-FLAG and FtsK-miniTurbo-FLAG were expressed from the *pilA* promoter, and sfGFP-miniTurbo-FLAG from the P_van_ in the presence of 100 µM vanillate added 18 h before the addition of 100 µM biotin and cephalexin for 4 h. Samples from three biological replicates were analysed. X-axis, log_2_-fold enrichment in experimental samples compared to sfGFP-miniTurbo-FLAG (negative control) calculated based on normalized intensities. Y-axis, -log_10_ of *p*-value. Significantly enriched proteins in the experimental samples (log_2_ ratio ≥3; *p-*value ≤0.01 (-log_10_ ≥ 2.0) are indicated by the stippled lines. DmxA and FtsK are shown in blue and red, respectively, and other enriched proteins in yellow. Enriched proteins other than FtsK and DmxA are listed in Table S1 and S2. **f-g**, Analysis of DmxA-mVenus variants. Domain architecture of variants (f) and population-based motility assay for T4P-dependent motility (g, upper panels) and localization (g, lower panels). Motility was analysed as in Fig. 2a. In lower panels, the percentage of cells with cluster at mid-cell is indicated. White arrowheads indicate clusters (n=200 from one biological replicate). Scale bars, 1 mm (upper panels), 5 μm (lower panels).

DmxA is encoded in an operon with the FtsB divisome protein, and this genetic organization is conserved in related species (Fig. S3a-d). However, Δ*dmxA* cells had neither a growth (Fig. S3e) nor a cell length defect (Fig. S3f). Therefore, to test whether DmxA is recruited to the division site by the divisome, we first treated cells with cephalexin, which blocks cytokinesis after initiation of constriction in *M. xanthus* ^45^. In cells treated with cephalexin for one generation, the frequency of cells with a constriction and a DmxA-mVenus cluster at mid-cell had significantly increased (Fig. 3a). DmxA-mVenus accumulated at the same level in cephalexin-treated and untreated cells (Fig. S4a), hinting that DmxA-mVenus synthesis is not cell cycle-regulated. Next, we depleted *dmxA-mVenus* expressing cells of the essential cell division protein FtsZ ^45^ using a strain in which the only copy of *ftsZ* was expressed from a vanillate-inducible promoter (P_van_). In the presence of vanillate, DmxA-mVenus localized at constrictions in 3% of cells (Fig. 3c). At 10 h of depletion, FtsZ was undetectable by immunoblotting (Fig. S4b), and cells had neither constrictions nor DmxA-mVenus clusters at mid-cell (Fig. 3c) despite the protein accumulating at a slightly higher level than in untreated cells (Fig. S4b). Finally, we observed in proximity labelling experiments using strains expressing either DmxA or sfGFP fused to the promiscuous biotin ligase miniTurbo (Fig. S4c) that the cell division protein FtsK, which localizes to the division site in *M. xanthus* ^46^, was significantly enriched in the DmxA-miniTurbo samples (Fig. 3d; Table S1). Equally, in the reciprocal experiment using an FtsK-miniTurbo construct (Fig. S4c; Table S2), DmxA was significantly enriched (Fig. 3e).

Altogether, these observations support that DmxA accumulation is not cell cycle-regulated and that DmxA interacts with protein(s) of the divisome, and is thereby recruited to the division site late during cytokinesis.

### DmxA function depends on DGC activity and is recruited to the division site by the TMD

To determine whether DGC activity and which domains contribute to DmxA function and localization, we replaced *dmxA* with *mVenus*-fused versions of full-length *dmxA*^WT^, *dmxA*^E626A^ and *dmxA*^R615A^ as well as with the truncated variants *dmxA*^ΔGAF×2^, *dmxA*^ΔTMD^ and *dmxA*^TMD^ (Fig. 3f). By immunoblot analysis, all variants accumulated at similar levels except DmxA^ΔGAF×2^-mVenus and *dmxA*^TMD^-mVenus, which accumulated at slightly lower levels (Fig. S5a). Among these variants, only DmxA^WT^-mVenus and DmxA^R615A^-mVenus supported WT motility (Fig. 3g; S5b). DmxA-mVenus localization to the division site was independent of DGC activity, the I-site and the two GAF domains (Fig. 3g). By contrast, the TMD was not only essential but also sufficient for DmxA-mVenus localization to the division site (Fig. 3g). Thus, DmxA function depends on DGC activity but not on c-di-GMP binding to the I-site, and DmxA is recruited to the division site by the TMD.

### DmxA DGC activity is activated upon recruitment to the division site

Based on the DmxA localization pattern, we hypothesized that DmxA activity is cell cycle-regulated and explicitly switched on late during cytokinesis. To test this idea, we used the genetically-encoded c-di-GMP biosensor cdGreen2 ^47^, for which binding of c-di-GMP results in conformational changes leading to increased green fluorescence, thus allowing real-time measurements of c-di-GMP levels at single-cell resolution over the entire cell cycle. For normalization of the cdGreen2 signal, *cdGreen2* was co-expressed with a gene encoding the fluorescent protein mScarlet-I ^47^.

For WT, we observed significant cell-to-cell variability of the cdGreen2 signal, while the mScarlet-I signal varied much less (Fig. 4a; S6a-b). Intriguingly, this cell-to-cell variability was clearly bimodal and only long cells with a constriction and some very short cells had a very high cdGreen2 signal (Fig. 4a; S6a). To focus on DmxA, we generated a mutant lacking 10 of the 11 predicted DGCs in *M. xanthus,* leaving DmxA as the only DGC (henceforth the Δ10 mutant). In the Δ10 mutant, DmxA-mVenus localized to the division site as in WT (Fig. S6c). Remarkably, the cdGreen2 signal was even more clearly bimodal in this strain, i.e. high in long cells with a constriction and in some very short cells (Fig. 4a; S6b). By contrast, all Δ*dmxA* cells, i.e. cells only lacking DmxA but retaining the remaining 10 DGCs, had the same low cdGreen2 background signal (Fig. 4a; S6b).

**Figure 4.**
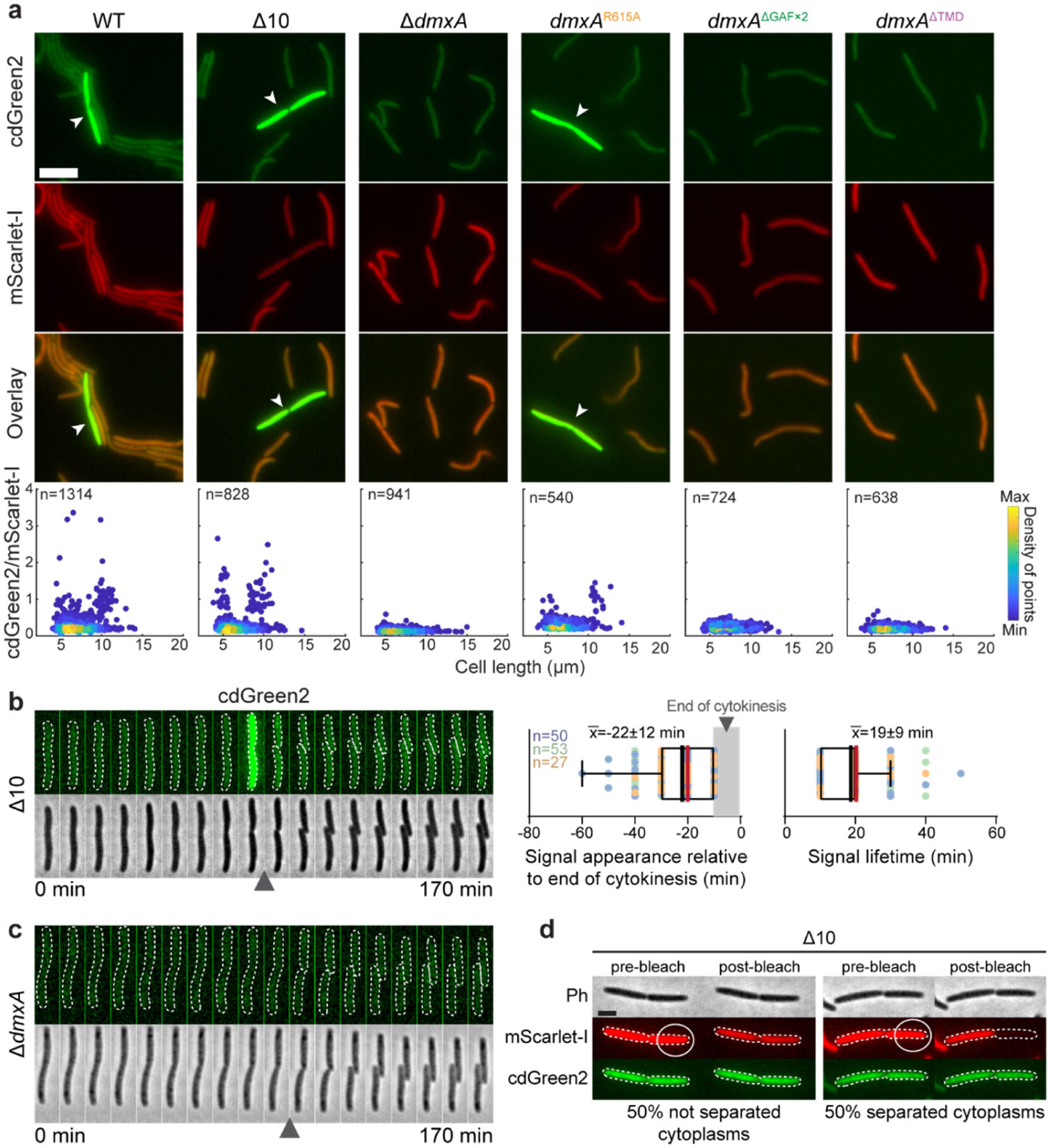
DmxA DGC activity is switched on upon recruitment to the division site. **a**, Analysis of cdGreen2 and mScarlet-I fluorescence in the indicated strains. Upper panels, epifluorescence snapshot images of cells expressing cdGreen2 and mScarlet-I. White arrowheads indicate cells with high cdGreen2 fluorescence. The *cdGreen2-mScarlet-I* operon was expressed from the constitutively active *pilA* promoter. Scale bar, 5 μm. Lower panels, scatter plots of the cdGreen2/mScarlet-I fluorescence ratio of each cell relative to its cell length. Colours indicate the density of points according to the scale on the right. Total number of cells from one biological replicate indicated. **b-c,** cdGreen2 fluorescence in Δ10 (b) and Δ*dmxA* (c) cells during the cell cycle. Left panels, epifluorescence and phase-contrast images from time-lapse microscopy of cells expressing cdGreen2. Images were recorded every 10 min; arrowheads indicate completion of cytokinesis. In b, right panels, analysis of the appearance relative to completion of cytokinesis and lifetime of the high cdGreen2 fluorescence. The first time point after completion of cytokinesis is defined as t=0 and indicated by the grey vertical bar. Box plots as in Fig. 3b based on three biological replicates with the number of cells per replicate and the corresponding data points in different colours. **d**, FRAP experiment on predivisional Δ10 cells expressing cdGreen2 and mScarlet-I. The mScarlet-I signal of one half of a cell was bleached. Post-bleached images were recorded 2 s after the bleaching event. All predivisional cells analysed had a high cdGreen2 fluorescent signal. n=22 from one biological replicate. Scale bar, 2 μm.

Time-lapse microscopy of Δ10 and WT cells revealed that the cdGreen2 signal increased dramatically shortly before completion of cytokinesis and then decreased rapidly in the two daughters, which had equal levels of cdGreen2 fluorescence (Fig. 4b; S6d). By contrast, Δ*dmxA* cells completely lacked the transient increase in c-di-GMP (Fig. 4c). The increase in c-di-GMP initiated 22±12 min before completion of cytokinesis, and remained high for 19±9 min (Fig. 4b). Remarkably, DmxA localizes to the division site with a similar timing (Fig. 3b). Moreover, in cephalexin-treated cells, the DmxA-mVenus clusters persisted longer at the division site and then eventually disintegrated (Fig. S6e). Notably, the high cdGreen2 signal also persisted longer in cephalexin-treated cells and eventually also vanished (Fig. S6f). We conclude that DmxA is sufficient and required for the transient c-di-GMP burst during cytokinesis.

The fully active DmxA^R615A^ I-site variant also supported the burst in c-di-GMP, while the non-complementing DmxA^ΔTMD^ and DmxA^ΔGAF×2^ variants did not (Fig. 4a; S6b). Importantly, DmxA^ΔTMD^-mVenus is similar to MalE-DmxA^WT^, which has DGC activity *in vitro* (Fig. 1d), but DmxA^ΔTMD^-mVenus does not localize to the division site (Fig. 4a), arguing that DmxA DGC activity *in vivo* is explicitly activated upon its recruitment to the division site. Furthermore, DmxA^ΔGAF×2^ localizes to mid-cell but lacks the region for DmxA dimerization, arguing that this region also *in vivo* mediates dimerization and, thus, DGC activity.

Finally, we observed no significant difference in the c-di-GMP levels in unsynchronized populations of WT and Δ*dmxA* cells (Fig. S6g), corroborating that DmxA only displays DGC activity for a brief period (∼20 min) of the 5 h cell cycle.

To resolve whether the increase in c-di-GMP initiates before or after the separation of the cytoplasm of the two daughters, we performed fluorescence recovery after photobleaching (FRAP) experiments in which we bleached the mScarlet-I signal in one half of pre-divisional cells with a high cdGreen2 signal. The bleaching event caused a decrease in the fluorescence signal in both cell halves in 50% of cells and only affected the signal in the bleached half in the remaining 50% (Fig. 4d).

We conclude that DmxA causes the transient c-di-GMP burst during cytokinesis and that DGC activity is explicitly activated upon recruitment to the division site and before the cytoplasm of the daughters is separated. By coupling DmxA activity and cytokinesis, two daughters with equal levels of c-di-GMP are generated.

### DmxA is essential for the symmetric incorporation and allocation of the core T4PM at the nascent and new cell poles

We next sought to establish the link between DmxA DGC activity during cytokinesis, motility and cell polarity. Because the T4PM core is present at both cell poles, is symmetrically incorporated at the nascent and new poles during and immediately after completion of cytokinesis ^10, 48^, and DmxA is active during cytokinesis, we initially focused on the polar assembly of the core T4PM. This assembly starts with forming the PilQ secretin in the outer membrane (OM), progresses inward, and culminates with the incorporation of cytoplasmic PilM ^10^ (Fig. S2b). The PilB and PilT ATPases bind to the cytoplasmic base of the core T4PM at the leading pole in a mutually exclusive manner to stimulate T4P extension and retraction, respectively ^9, 49, 50^ (Fig. S2b). Note that PilT localizes bipolarly, while PilB almost exclusively localizes unipolarly to the leading pole ^9, 21^.

First, we performed time-lapse microscopy using PilQ-sfGFP as a readout for assembly of the core T4PM. As reported, in WT, PilQ-sfGFP incorporation was initiated during cytokinesis and, upon completion of cytokinesis, continued symmetrically at the new poles in the two daughters (Fig. 5a; S7), giving rise to mirror-symmetric daughters. Strikingly, in the absence of DmxA, PilQ-sfGFP incorporation was not only significantly delayed but also asymmetric in the two daughters, resulting in asymmetric daughter pairs and with many cells having PilQ-sfGFP clusters of very different intensities or even only a single, unipolar PilQ-sfGFP cluster at the old pole (Fig. 5a; S7). Moreover, this defect in PilQ-sfGFP polarity was typically not fully corrected before the subsequent cell division (Fig. 5a; S7), resulting in propagation of the polarity defect.

**Figure 5.**
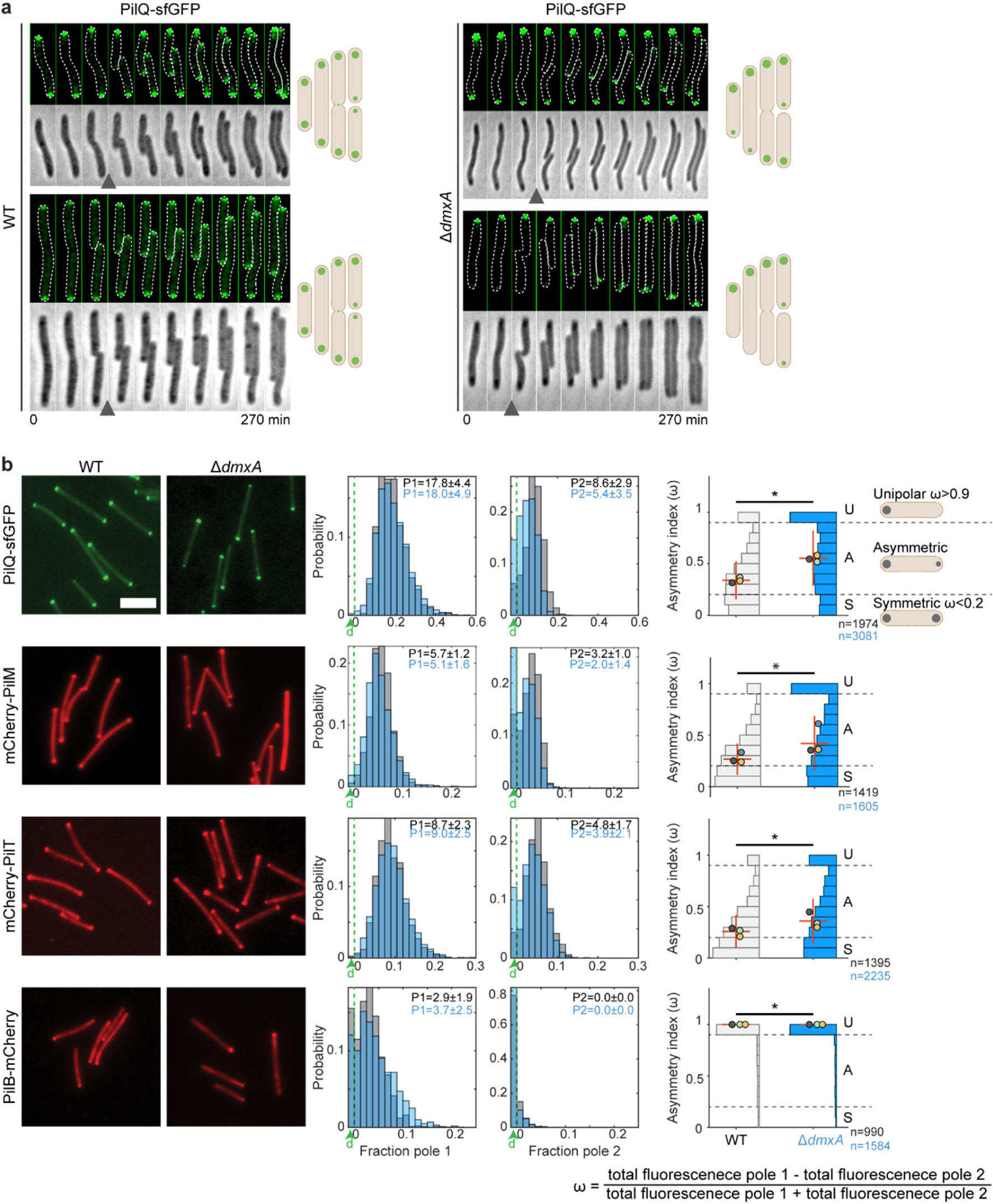
DmxA is essential for the symmetric incorporation and allocation of the core T4PM proteins at the nascent and new cell poles. **a**, PilQ-sfGFP localization during the cell cycle. Epifluorescence and phase-contrast images from time-lapse microscopy of WT (left panels) and Δ*dmxA* cells (right panels). Images were recorded every 30 min; arrowheads indicate completion of cytokinesis. Schematics show dominant localization patterns of PilQ-sfGFP in WT and Δ*dmxA* cells during the cell cycle. **b**, Localization of PilQ-sfGFP, mCherry-PilM, PilB-mCherry and mCherry-PilT in WT and Δ*dmxA* cells. Left panels, representative snapshot images. Scale bar, 5 μm. Middle panels, histograms of the distribution of the fraction of the total cellular fluorescence in polar clusters at pole 1 and pole 2 in WT (grey) and Δ*dmxA* (blue) cells. The pole with highest fluorescence is defined as Pole 1. Numbers in the upper right corners indicate the median ± MAD fluorescence signal at pole 1 (P1) and pole 2 (P2). The fraction of cells with no polar signal(s) is indicated in the leftmost column labelled d (for diffused) in green; cells in which no polar cluster was detected at pole 1 also do not have a signal at pole 2. Right panels, histograms of the distribution of the asymmetry index (ω). Localization patterns are binned as unipolar, asymmetric and symmetric from the ω values as indicated; cells in which no polar signal was detected were not considered in the analysis. Error bars indicate median ± MAD. Differently coloured circles indicate the median of each of three biological replicates. The total number of analysed cells is indicated below. **\****p*<0.05, Mann-Whitney test.

Consistent with the faulty polar PilQ incorporation, we observed in quantitative analyses of snapshot microscopy images that PilQ-sfGFP, mCherry-PilM and mCherry-PilT localization in the Δ*dmxA* mutant was significantly shifted toward unipolar and, thus less symmetric than in WT (Fig. 5b). Similarly, PilB-mCherry was significantly more unipolar in Δ*dmxA* cells. The shifts toward asymmetry were primarily caused by many Δ*dmxA* cells having no or a strongly decreased fluorescence signal at the pole with the lowest fluorescence (Fig. 5b). Thus, in the absence of DmxA, all tested T4PM proteins localize significantly more asymmetrically than in WT.

We conclude that DmxA and the burst in c-di-GMP are essential for the symmetric incorporation and allocation of PilQ during and after cytokinesis, thereby generating mirror-symmetric daughters. Moreover, we infer that the faulty polar PilQ incorporation and allocation contribute to the more asymmetric localization of the other tested T4PM proteins in the Δ*dmxA* mutant.

### DmxA is essential for the symmetric allocation of the polarity proteins to the daughters during cytokinesis

We predicted that the proteins of the polarity module would also be symmetrically allocated during cytokinesis to generate mirror-symmetry of these proteins in the daughters. Briefly, RomR alone localizes polarly in the absence of the remaining five polarity proteins and brings about polar localization of these proteins ^29, 32^. Moreover, RomX and MglC localization follows that of RomR ^26, 29^ and RomY the highest concentration of MglB ^27^ (Fig. S2f). Therefore, we used RomR and MglB as well as MglA, which generates the output of the polarity module, as readouts for the localization of the proteins of the polarity module.

First, to test our prediction, we performed time-lapse microscopy of WT expressing RomR-mCherry. These analyses revealed a precise order of events in which, late during cytokinesis, RomR-mCherry was released from the old poles, then localized to the division site, and, upon completion of cytokinesis, was symmetrically allocated to the two daughters (Fig. 6a), giving rise to mirror-symmetric daughters. Remarkably, in the absence of DmxA, we observed very different patterns. During most division events, RomR-mCherry was released from the old poles, however, it was either not recruited at the division site but instead switched to the opposite pole (Fig. 6a, upper), or if it localized to the division site, it was asymmetrically allocated to the daughters (Fig. 6a, lower). Consequently, the daughters of a division event contained different amounts of RomR-mCherry, were not mirror-symmetric, and had polar clusters of very different intensities or even only a single RomR-mCherry cluster at the old pole. The defects in RomR-mCherry polarity were typically also not fully corrected before the subsequent cell division (Fig. 6a).

**Figure 6.**
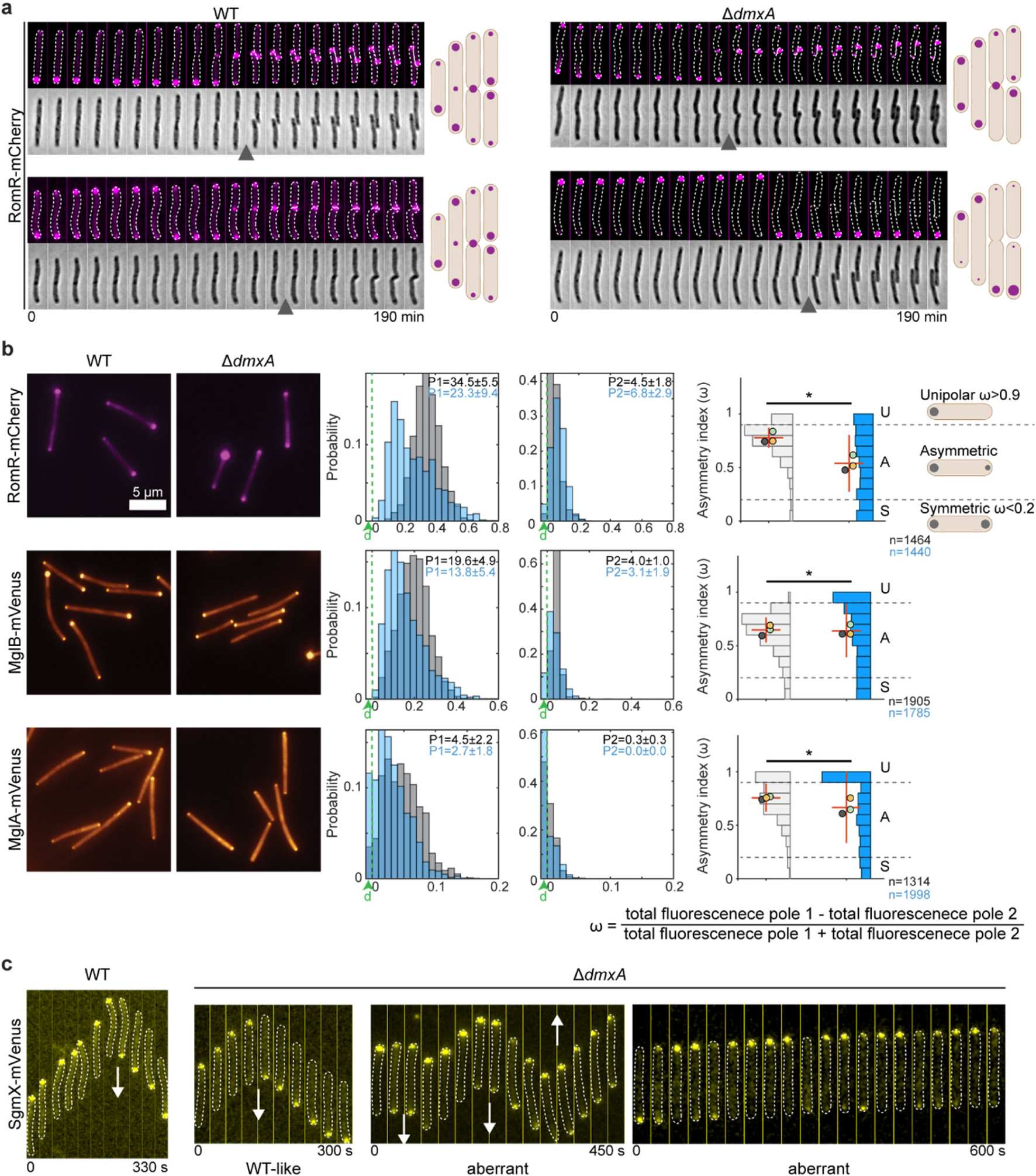
DmxA is essential for the symmetric allocation of polarity proteins to the daughters during cytokinesis. **a**, RomR-mCherry localization during the cell cycle. Epifluorescence and phase-contrast images from time-lapse microscopy of WT and Δ*dmxA* cells. Images were recorded every 10 min; black arrowheads indicate completion of cytokinesis. Schematics show localization patterns of RomR-mCherry in WT and Δ*dmxA* cells during the cell cycle. **b**, Localization of RomR-mCherry, MglB-mVenus, MglA-mVenus in WT and Δ*dmxA* cells. Left panels, representative snapshot images. Analysis of snapshot images of WT (grey) and Δ*dmxA* (blue) were done as in Fig. 5b. **c**, SgmX-mVenus localization in moving WT and Δ*dmxA* cells. Images were recorded every 30 s. White arrows indicate reversals.

Consistent with the faulty polar RomR allocation to the daughters in the absence of DmxA, we observed in quantitative analyses of snapshot microscopy images that RomR-mCherry, MglB-mVenus and MglA-mVenus in the Δ*dmxA* mutant localized in highly aberrant patterns and had largely lost their defined polar asymmetry in individual cells, displaying much broader variations in asymmetry values compared to WT (Fig. 6b). The aberrant RomR-mCherry and MglB-mVenus asymmetry resulted from the much broader variations in the fluorescence signals at both poles (Fig. 6b); accordingly, MglA polar localization was significantly reduced in many cells (Fig. 6b).

We conclude that RomR polarity is reset during cytokinesis in WT and that DmxA together with the burst in c-di-GMP are essential for this reset and the symmetric allocation of RomR to the two daughters. Moreover, we infer that the highly aberrant localization of MglB and MglA in the Δ*dmxA* mutant results from the faulty RomR allocation during cytokinesis.

Finally, we aimed to establish the link between the aberrant localization of the T4PM and the polarity proteins to the aberrant T4P-dependent motility behaviour of the Δ*dmxA* cells. Because the polar signals of PilB-mCherry and MglA-mVenus are low, it is technically difficult to follow these fusions in time-lapse recordings. Therefore, we followed the localization of the MglA-GTP effector SgmX-mVenus, which is recruited to the leading pole by MglA, and then recruits the PilB ATPase to stimulate T4P extension ^20, 21^. In WT, SgmX-mVenus localized with a large cluster at the leading pole in 100% of cells and switched polarity during reversals (n=48) (Fig. 6c). In the Δ*dmxA* mutant, SgmX-mVenus localized as in WT in ∼75% of cells (n=84) but in the remaining ∼25%, SgmX-mVenus localized aberrantly with either a bipolar pattern and/or more unstably at the leading pole, i.e. the intensity at the leading pole would shortly decrease, and this was occasionally accompanied by a brief increase in fluorescence at the opposite pole (Fig. 6c). Importantly, many of these ∼25% of cells hyper-reversed (Fig. 6c).

Altogether, these observations support that the aberrant localization of the T4PM and the polarity proteins caused by lack of DmxA results in motility defects with aberrant reversals.

## Discussion

Here, we describe a c-di-GMP-dependent genetic program that is hardwired into the *M. xanthus* cell cycle and guarantees the formation of mirror-symmetric, phenotypically similar daughter cells. Specifically, the DGC DmxA is explicitly recruited to the division site late during cytokinesis, and released upon completion of cytokinesis. During this brief period of the cell cycle, its DGC activity is switched on resulting in a dramatic but transient increase in the c-di-GMP concentration. This c-di-GMP burst, in turn, ensures the equal and symmetric allocation of core T4PM proteins and polarity proteins to the two daughters. In the absence of DmxA, the daughters inherit unequal amounts of these proteins causing aberrant T4PM localization and cell polarity and, consequently, aberrant motility behaviour. Thus, *M. xanthus* harnesses DmxA and c-di-GMP to ensure the generation of mirror-symmetric, phenotypically similar daughters in each cell division event.

DmxA recruitment to the division site late during cytokinesis depends on the TMD of DmxA and the divisome, suggesting that the TMD interacts with the divisome. Indeed, using proximity labelling, we identified the transmembrane divisome protein FtsK as a potential direct interaction partner of DmxA. Several lines of evidence support that DmxA is activated upon recruitment to the division site. First, the timing of DmxA localization to the division site and the burst in c-di-GMP perfectly correlate. Second, DmxA is required and sufficient for the burst in c-di-GMP. Third, DmxA^ΔTMD^ does not localize to the division site and is not active *in vivo*; however, the protein has DGC activity *in vitro*. Although we cannot rule out that DmxA^ΔTMD^ is less active than full-length DmxA, these observations jointly support that DmxA is activated at the division site. Finally, in cephalexin-treated cells and in cells depleted of FtsZ, DmxA accumulates at the same level as in untreated cells, strongly indicating that DmxA accumulation is not cell cycle-regulated and that DmxA activity is not regulated by the total cellular concentration. Based on these observations, we suggest that DmxA DGC activity, upon recruitment to the division site, is switched on either by interacting with protein(s) of the divisome or, alternatively, the high local DmxA concentration stimulates formation of the enzymatically active dimer. We speculate that the low-affinity I-site allows DmxA to synthesize high concentrations of c-di-GMP, and may solely be relevant at very high concentrations to avoid excessive overproduction of c-di-GMP. Upon completion of cytokinesis, the DmxA cluster disintegrates, likely subsequent to the disassembly of the divisome. As a consequence, c-di-GMP synthesis ceases and its level decreases rapidly. Interestingly, none of the six predicted PDEs of *M. xanthus* have been implicated in motility^34^. In the future, it will be interesting to determine which PDE(s) are involved in the rapid decrease in the c-di-GMP concentration.

Despite only being active during a brief period of the cell cycle, DmxA is essential for WT motility behaviour. During this brief period, DmxA guarantees the symmetric incorporation and allocation at the nascent and new poles of PilQ of the T4PM and RomR of the polarity module. Because PilQ and RomR are at the base of the assembly of the core T4PM and the polar localization of the proteins of the polarity module, respectively, we suggest that the defects in PilQ and RomR polar localization during cytokinesis in the absence of DmxA causes the observed misincorporation of the core T4PM and the remaining polarity regulators, respectively. Because the polarity defects that arise during one division in cells lacking DmxA are not fully corrected until the next division, and mutants with aberrant localization of the polarity proteins have aberrant cell behaviours with altered reversal frequencies ^18, 23, 24, 26, 27, 29, 33^, we suggest that the aberrant T4P-dependent motility behaviour in the absence of DmxA is the result of dual defects, i.e. the defects in the polar incorporation of the core T4PM and in the localization of polarity proteins. We also speculate, but have not shown, that the aberrant gliding behaviour is not only a consequence of the mislocalized polarity proteins but also involves defects in the polar incorporation of structural proteins of the Agl/Glt machine.

How, then, does the burst in c-di-GMP ensure the correct incorporation and allocation of polarly localized motility proteins and regulators? The effects of changing c-di-GMP levels are implemented by the binding of c-di-GMP to downstream effectors ^2, 3^. Because polar PilQ incorporation depends on its peptidoglycan-binding AMIN domains ^51^, we suggest that the c-di-GMP burst brings about the localization of a landmark protein, which possibly binds peptidoglycan, at the nascent and new poles that assist polar recruitment of PilQ. The cytoplasmic RomR protein by an unknown mechanism localizes polarly in the absence of the other polarity proteins ^32^. The symmetric allocation of RomR to the daughters involves a polarity reset involving three steps, i.e. RomR release from the old poles, its recruitment to the division site, and its symmetric allocation to the two daughters. In the absence of DmxA, RomR was still released from the old poles in most cells, suggesting that this step is independent of DmxA and c-di-GMP. However, RomR was either not recruited to the division site or, if it was recruited, then it was not symmetrically allocated to the two daughters. Therefore, we suggest that c-di-GMP brings about the localization of a landmark protein at the division site that is recognized by RomR. Because the defects in PilQ and RomR polar localization are not fully corrected between division events, we also suggest that these landmarks may only be transiently active. In future experiments, it will be important to identify the effector(s) involved in the response to DmxA-generated c-di-GMP and to address whether these effector(s) serve as landmark(s) or to recruit landmark(s). Nonetheless, we speculate that an advantage of engaging a DGC in setting up correct cell polarity during cytokinesis could be that c-di-GMP would allow the transient function of effector(s)/polar landmark(s).

Several bacteria that alternate between a planktonic, flagellum-dependent swimming lifestyle and a surface-associated lifestyle, harness c-di-GMP to deterministically generate phenotypically distinct daughters during division ^4, 5^. In *Caulobacter crescentus*, *Pseudomonas aeruginosa* and *Shewanella putrefaciens*, the genetic programs driving the generation of this heterogeneity rely on the asymmetric deployment of c-di-GMP metabolizing enzyme(s) to the daughters during cell division, i.e. either the relevant DGC and PDE localize to opposite cell poles or a PDE localizes unipolarly^40, 47, 52–59^ (Fig. 7). Consequently, one daughter has low c-di-GMP and becomes the flagellated swimming daughter, while the other daughter has high c-di-GMP and becomes the surface-associated daughter. By contrast, *M. xanthus* places the DGC DmxA at the division site, thereby enabling the formation of mirror-symmetric and phenotypically similar daughters. Thus, by deploying c-di-GMP synthesizing and degrading enzymes to distinct subcellular locations, bacteria harness c-di-GMP to establish deterministic genetic programs that are hardwired into the cell cycle to generate or, as shown here, minimize phenotypic heterogeneity (Fig. 7).

**Figure 7.**
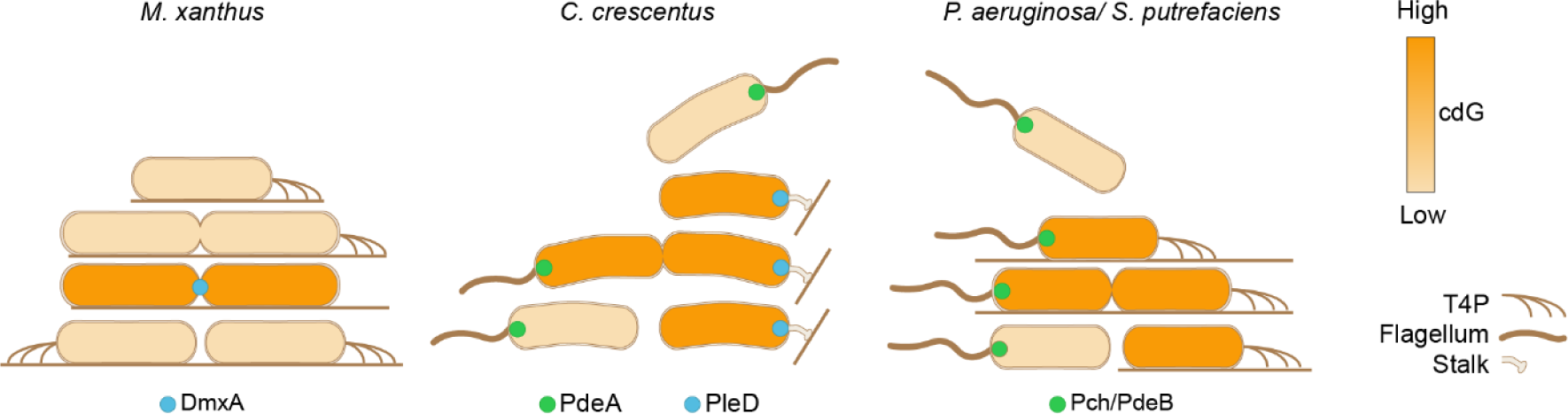
Deployment of DGC and/or PDEs to distinct subcellular locations establishes deterministic genetic programs hardwired into the cell cycle to generate or minimize phenotypic heterogeneity. In *M. xanthus,* DmxA (blue) localizes to and is switched on at the division site creating a c-di-GMP burst that ensures similar daughter cells. In *C. crescentus,* the flagellated, swarmer cell has low c-di-GMP due to the activity of the PDE PdeA at the flagellated pole (green). Upon differentiation to the surface-associated stalked cell, the c-di-GMP level increases due to the activity of the DGC PleD at the stalked pole (blue). In predivisional cells, PdeA and PleD localize to opposite poles, giving rise to a swarmer cell with low c-di-GMP and a stalked cell with high c-di-GMP upon division. In *P. aeruginosa*/*S. putrefaciens,* the flagellated, swimming cell has low c-di-GMP due to the activity of the PDE Pch/PdeB at the flagellated pole (green). Upon surface contact, c-di-GMP increases but the involved DGC(s) remain to be identified. High c-di-GMP stimulates T4P formation and surface adhesion. During division, the flagellated pole inherits the PDE, creating a flagellated, swimming daughter with low c-di-GMP and a surface-adhered, piliated daughter with high c-di-GMP.

Similar to stochastically generated phenotypic heterogeneity ^60–62^, the deterministic generation of phenotypic heterogeneity has been suggested to be part of a bet-hedging strategy in which the diversification of phenotypes optimizes the survival of the population and/or a division of labour strategy by enabling the colonization of multiple habitats in parallel ^55, 58, 59, 63^. Because *M. xanthus* translocates on surfaces in large cooperative swarms in which the motility of individual cells is highly coordinated, we speculate that reducing phenotypic heterogeneity during cell division optimizes its ability to perform its motility-dependent social behaviours.

## Acknowledgement

We thank Sean Murray for helpful discussions and Dorota Skotnicka, Anke Treuner-Lange and Memduha Muratoglu for strains, and the Research Core Unit Metabolomics at the Hannover Medical School for the assistance with measuring c-di-GMP levels. This work was supported by Deutsche Forschungsgemeinschaft (DFG, German Research Council) within the framework of the SFB987 “Microbial Diversity in Environmental Signal Response”, by the Max Planck Society, and by the Swiss National Science Foundation grant 310030_208107.

## Availability of data and materials

The authors declare that all data supporting this study are available within the article and its Supplementary Information files. All materials used in the study are available from the corresponding author.

## Methods

### Bacterial strains and growth media

*M. xanthus* cells were grown at 32°C in 1% CTT (1% (w/v) Bacto Casitone (Gibco) in TPM buffer (10 mM Tris-HCl pH 8.0, 1 mM K_2_HPO_4_/KH_2_PO_4_ pH 7.6, and 8 mM MgSO_4_)) liquid medium or on 1.5% agar supplemented with 1% CTT ^64^. Oxytetracyline and kanamycin at concentrations of 10 μg ml^−1^ and 50 µg ml^−1^, respectively were added when needed. Cephalexin was added to a concentration of 35 µg ml^−1^ in liquid medium and 20 µg ml^−1^ on agarose. All *M. xanthus* strains are derivatives of the WT strain DK1622 ^12^. *M. xanthus* strains, plasmids and oligonucleotides used in this work are listed in Tables S3-S5, respectively. In-frame deletions or gene replacements were generated as described ^65^, plasmids were integrated in a single copy by site-specific recombination at the Mx8 *attB* site or by homologous recombination at the *MXAN_18-19* site or at the endogenous locus. All in-frame deletions and plasmid integrations were verified by PCR. Plasmids were propagated in *E. coli* Mach1, which was grown at 37°C in lysogeny broth (LB) medium (10 mg tryptone ml^−1^, 5 mg yeast extract ml^−1^ and 10 mg NaCl ml^−1^) supplemented when required with kanamycin (50 µg ml^−1^) or tetracycline (25 µg ml^−1^).

### Motility assays

Population-based motility assays were performed as described ^66^. Briefly, exponentially growing suspension cultures were harvested (3 min, 8,000 *g*, RT) and resuspended in 1% CTT to a calculated density of 7×10^9^ cells ml^−1^. 5 µl aliquots of cell suspensions were spotted on 0.5% agar (Invitrogen) and 1.5% agar (Gibco) supplemented with 0.5% CTT and incubated at 32°C. Cells were imaged at 24 h using a M205FA Stereomicroscope (Leica) and a DMi8 inverted microscope (Leica) equipped with a Hamamatsu ORCA-Flash4.0 V2 digital CMOS C11440 camera (Hamamatsu Photonics) and DFC9000 GT camera (Leica), respectively. To visualise single cells moving by T4P-dependent motility, 5 µl exponentially growing cells in suspension were placed in a 24-well polystyrene plate (Falcon). After 10 min incubation in the dark at RT, 200 µl of 1% methylcellulose in MMC buffer (10 mM MOPS, 4 mM MgSO_4_, 2 mM CaCl_2_, pH 7.6) were added, and cells incubated for 30 min in the dark at RT. Cells were imaged for 15 min with 30 s intervals. To visualize individual cells moving by gliding, exponentially growing cells in suspension were diluted to 3×10^9^ and 5 µl spotted on 1.5% agar (Gibco) supplemented with 0.5% CTT and immediately covered with a coverslide. Cells were incubated 2 h at 32°C and then visualized for 15 min with 30 s intervals at RT. Cells were imaged using a DMi8 Inverted microscope and DFC9000 GT camera. Images were analysed using ImageJ ^67^.

### Negative stain transmission electron microscopy

10 µl of *M. xanthus* cells exponentially grown in suspension were placed on one side of the electron microscopy grid (Plano) and incubated at RT for 40 min. To avoid evaporation during this step, the grid was incubated in humid air. The liquid was blotted through the grid by capillarity by applying the side of the grid on Whatman paper. Cells were washed three times with 10 µl of double-distilled water and stained with UA-Zero EM Stain (Plano) (diluted to 0.25% (v/v) in double distilled water). After 1 min incubation, the liquid was removed and cells washed once with double-distilled water to remove excess staining solution. Transmission electron microscopy was done with a JEOL JEM-1400 electron microscope at 100 kV.

### Immunoblot analysis

Immunoblots were performed as described ^68^. Rabbit polyclonal α-LonD (dilution: 1:5000) ^69^, α-PilC (dilution: 1:2,000) ^9^, α-FtsZ (dilution: 1:25,000) ^45^, α-mCherry (dilution: 1:1000) (BioVision) and α-RFP (dilution 1:2,000) (Rockland), were used together with horseradish peroxidase-conjugated goat α-rabbit immunoglobulin G (dilution: 1:15,000) (Sigma) as secondary antibody. Mouse α-GFP antibodies (dilution: 1:2,000) (Roche) were used together with horseradish peroxidase-conjugated sheep α-mouse immunoglobulin G (dilution: 1:2000) (GE Healthcare) as a secondary antibody. Blots were developed using Luminata Forte Western HRP Substrate (Millipore) on a LAS-4000 imager (Fujifilm).

### Operon mapping

Mapping of the *ftsB-dmxA* operon was performed as described ^51^. Briefly, 1×10^9^ WT cells from an exponentially growing suspension culture were harvested (3 min, 8,000 *g,* room temperature (RT)) and resuspended in 200 μl lysis-buffer (100 mM Tris-HCl pH 7.6, 1 mg ml^−1^ lysozyme). After incubation at 25°C for 5 min, cells were lysed and RNA purified using the Monarch Total RNA Miniprep Kit (NEB) according to the manufacturer’s instructions except that the on-column DNase treatment was omitted. RNA was eluted in RNase-free water, treated with Turbo DNase (Invitrogen) and purified using the Monarch RNA Cleanup Kit (50 μg) (NEB) and eluted in RNase-free water. 1 μg of RNA was used for cDNA synthesis using the LunaScript RT SuperMix Kit (NEB) with and without reverse-transcriptase. cDNA, the mock reaction without reverse-transcriptase, or genomic DNA were used as template for PCR using the primers listed in Table S5.

### Cell length determination

5-µl aliquots of exponentially growing suspension cultures were spotted on 1% agarose supplemented with 0.2% CTT. Cells were immediately covered with a coverslide, and imaged using a DMi8 Inverted microscope and DFC9000 GT camera. To assess cell length, cells were segmented using Omnipose ^70^, segmentation was manually curated using Oufti ^71^, analysed using Matlab R2020a (The MathWorks) and plotted using GraphPad Prism (GraphPad Software, LLC).

### Fluorescence microscopy

In all time-lapse microscopy experiments except for those involving SgmX-mVenus, cells were visualized as in ^72^ with slight modifications. Briefly, 5 µl exponentially growing cells in suspension were placed on a glass coverslide attached to a plastic frame. Cells were covered with a thick 1% agarose pad supplemented with 0.2% CTT, the pad sealed with parafilm to reduce evaporation, and cells imaged after 180 min. To avoid that cells would move out of the field of view, strains contained the Δ*gltB* mutation. To clearly distinguish leading and lagging cell poles of cells moving by T4P-dependent motility, time-lapse microscopy experiments involving SgmX-mVenus, were done on Chitosan-coated µ-Dishes (Ibidi) as described ^73^. Briefly, a 100 µl-aliquot of exponentially growing cells was diluted in 900 µl MC7 buffer (10 mM MOPS pH 7.0, 1 mM CaCl_2_), spotted on the chitosan-coated µ-Dish, and imaged after 30 min. Snapshot microscopy images were captured from cells on 1% agarose pad supplemented with 0.2% CTT (biosensor and DmxA-mVenus) or on Chitosan-coated µ-Dishes (all other fluorescent proteins). Cells were imaged using a DMi8 inverted microscope and a Hamamatsu ORCA-Flash4.0 V2 Digital CMOS C11440 or a DFC9000 GT camera. Data was analysed using Oufti ^71^, Metamorph® v 7.5 (Molecular Devices), Matlab and ImageJ ^67^. DmxA-mVenus clusters and constrictions were identified manually.

To identify and analyse polar clusters in snapshots, we used a custom-made Matlab script^26^. Briefly, cells were segmented, and polar clusters were identified as having an average fluorescence signal of 1.5 SD (MglA) or 2 SD (all other proteins), above the mean cytoplasmic fluorescence and a size of three or more pixels. For each cell with polar clusters, an asymmetry index (ω) was calculated as:

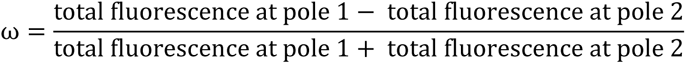

Pole 1 was assigned to the pole with the highest fluorescence. The localization patterns were binned from the ω values as follows: unipolar (ω>0.9), bipolar asymmetric (0.9≥ω>0.2) and bipolar symmetric (0.2≥ω). Diffuse localization was determined when no polar signal was detected. Polar fluorescence of moving cells synthesizing SgmX-mVenus was followed manually.

For the analysis of single cell cdGreen2 fluorescence, cells were segmented using Omnipose ^70^ and the segmentation manually curated using Oufti ^71^. For normalization, the average cellular fluorescence of each cell in the green channel (cdGreen2) was divided by the red channel (mScarlet-I).

### C-di-GMP quantification

C-di-GMP levels were determined as described ^74^. Briefly, 4 ml of exponentially growing cells were harvested by centrifugation (20 min, 2,500 *g*, 4°C). Cells were mixed with 300 µl ice-cold extraction buffer (high-pressure liquid chromatography [HPLC]-grade acetonitrile-methanol-water [2:2:1, v:v:v]), and incubated 15 min at 4°C to quench metabolism. Extraction was performed at 95°C for 10 min, samples were centrifuged (10 min, 21,130 *g*, 4°C) and the supernatant containing extracted metabolites transferred to a new Eppendorf tube. The pellet was washed with 200 µl extraction buffer and centrifuged (10 min, 21,130 *g*, 4°C). This step was repeated. The residual pellet containing proteins was kept, and the three supernatants from the extraction and the two washing steps containing c-di-GMP were pooled and evaporated to dryness in a vacuum centrifuge. Subsequently, the samples with c-di-GMP were dissolved in HPLC-grade water for analysis by liquid chromatography-coupled tandem mass spectrometry (LC-MS/MS). In parallel, to determine the protein concentration for each sample, the residual pellets were resuspended in 800 µl 0.1 M NaOH, and heated for 15 min at 95°C until dissolution. Protein levels were determined using a 660 nm Protein Assay (Pierce) following the manufacturer’s instructions.

### Protein purification

For expression and purification of MalE-tagged DmxA variants, proteins were expressed in *E. coli* Rosetta DE3 growing in 5052-Terrific-Broth ^75^ (0.5% (v/v) glycerol, 0.05% (w/v) glucose, 0.2% (w/v) lactose, 2.4% (w/v) yeast extract, 2% (w/v) tryptone, 25 mM Na_2_HPO_4_, 25 mM KH_2_PO_4_, 50 mM NH_4_Cl, 5 mM Na_2_SO_4_, 2 mM MgSO_4_) auto-induction medium supplemented with 25 µg ml^−1^ chloramphenicol and 100 µg ml^−1^ carbenicillin. Cells were grown at 37°C until OD_600_=1, shifted to 18°C and further incubated overnight. Cells were harvested and resuspended in MalE-lysis buffer (100 mM Tris-HCl pH7.2, 500 mM NaCl, 10 mM MgCl_2_, 5 mM DTT) supplemented with EDTA-free protease inhibitor cocktail (Roche) and lysed by sonication for ten cycles of 30 pulses of sonication and 30 s breaks using a Hielscher UP200st set to pulse=70%, amplitude=70%. The lysate was cleared by centrifugation (16,000 *g,* 4°C, 30 min) and loaded onto a 5 ml HighTrap MBP column (Cytiva) using an Äkta-Pure system (GE Healthcare). The column was washed with 10 column volumes of lysis buffer and protein eluted with MalE-elution buffer (100 mM Tris-HCl pH7.2, 500 mM NaCl, 10 mM MgCl_2_, 5 mM DTT, 10 mM Maltose). The elution fractions containing MalE-DmxA variants were pooled and loaded on a HiLoad 16/600 Superdex 200pg (GE Healthcare) SEC column, which was pre-equilibrated with SEC-buffer (50 mM Tris-HCl pH7.2, 250 mM NaCl, 10 mM MgCl_2_, 5 mM DTT, 5% glycerol (v/v)) and protein was eluted using SEC-buffer. Subsequently, protein was either used fresh or snap-frozen in SEC-buffer.

### DGC activity assay

DGC activity assays were performed, using the EnzCheck®Pyrophosphate Assay Kit (Thermo) as described ^76^. Briefly, the release of inorganic pyrophosphate during c-di-GMP synthesis was followed by measuring the absorbance change at 360 nm in a Tecan M200 pro, in 30 s intervals for 1 h. Reactions contained 1 µM protein, and 50 µM GTP. Inhibition reactions were fit in GraphPad Prism to the equation V_[cdG]_=V_0_/(1/1+([cdG]/K_i_)^h^), where V_0_ represents the reaction velocity in the absence of c-di-GMP, [cdG] the concentration of c-di-GMP in the reaction, K_i_ the inhibitory constant and h the Hill coefficient.

### *In vitro* nucleotide binding assay

C-di-GMP binding was determined by Bio-Layer Interferometry using the BLItz system (ForteBio) ^77^ and a Streptavidin SA biosensor (ForteBio). Briefly, 500 nM biotinylated c-di-GMP (Biolog) in SEC-buffer supplemented with 0.1% (v/v) Tween-20 was immobilized onto the biosensors for 120 s, and unbound molecules washed off for 30 s. Association and dissociation of a protein were carried out for 120 and 120 s respectively. Binding was fitted to the “One site – Total” binding model in GraphPad Prism.

### Proximity labelling

Proximity labelling including shotgun proteomics analysis was done as described ^78^. Briefly, 50 ml of exponentially growing cell suspension were incubated with 100 µM biotin, and 35 µg ml^−1^ cephalexin. After 4 h, cells were harvested by centrifugation (8,000 *g,* 10 min, 4°C), resuspended in 600 µl RIPA buffer (50 mM Tris-HCl pH 7.0, 150 mM NaCl, 0.5% (w/v) sodium deoxycholate, 0.2% (w/v) SDS, 1% (v/v) Triton-X100) supplemented with protease inhibitor cocktail (Roche) and lysed by 30 pulses of sonication using a Hielscher UP200st set to pulse 50%, amplitude 50%. SpinTrap G-25 columns (Cytiva) were used to remove an excess of biotin from the cleared lysate. To enrich biotinylated proteins, 500 µl of each sample was incubated for 1 h at 4°C with 50 µl streptavidin magnetic beads (Pierce). The beads were washed three times with 1 ml RIPA buffer, twice with 1 ml 1 M KCl, and three times with 1 ml 50 mM Tris-HCl pH 7.6. Finally, proteins were eluted using on-bead digest as described ^69^. Briefly, 100 µl elution buffer 1 (100 mM ammonium bicarbonate, 1 µg trypsin (Promega)) was added to each sample. After 30 min incubation at 30°C, the supernatant containing the digested proteins was collected. Beads were washed twice with elution buffer 2 (10 mM ammonium bicarbonate, 5 mM Tris(2-carboxyethyl)phosphine hydrochloride (TCEP)) and added to the first elution fraction. Digestion continued overnight at 30°C. Next, the peptides were incubated with 10 mM iodoacetamide for 30 min at 25°C in the dark. Prior to LC-MS analysis, peptide samples were desalted using C18 solid phase extraction spin columns (Macherey-Nagel). Peptide mixtures were then analysed using LC-MS on an Exploris 480 instrument connected to an Ultimate 3000 RSLCnano and a nanospray flex ion source (all Thermo Scientific). A detailed description of the LC-MS parameters are described in ^78^. The following separating gradient was used: 98% solvent A (0.15% formic acid) and 2% solvent B (99.85% acetonitrile, 0.15% formic acid) to 30% solvent B over 40 min at a flow rate of 300 nl/min. Peptides were ionized at a spray voltage of 2.3 kV, and ion transfer tube temperature set at 275°C, 445.12003 m/z was used as internal calibrant. The data acquisition mode was set to obtain one high-resolution MS scan at a resolution of 60,000 full width at half maximum (at m/z 200) followed by MS/MS scans of the most intense ions within 1 s (cycle 1s). The ion accumulation time was set to 50 ms (MS) and 50 ms at 17,500 resolution (MS/MS). The automatic gain control (AGC) was set to 3×10^6^ for MS survey scan and 2×10^5^ for MS/MS scans. MS raw data was then analysed with MaxQuant ^79^, and an *M. xanthus* UniProt database ^80^. MaxQuant was executed in standard settings without “match between runs” option. The search criteria were set as follows: full tryptic specificity was required (cleavage after lysine or arginine residues); two missed cleavages were allowed; carbamidomethylation (C) was set as fixed modification; oxidation (M) and deamidation (N,Q) as variable modifications. The MaxQuant proteinGroups.txt file was further processed by the SafeQuant R package for statistical analysis ^81^.

### Total proteome analysis

The total proteome of *M. xanthus* cells grown in suspension culture was determined following a slightly modified protocol of ^69^. Briefly, 2 ml of exponentially growing suspension cultures were harvested (8,000 *g,* 3 min, RT). Cells were resuspended in 1 ml PBS and harvested again. Subsequently, the supernatant was discarded and the pellet snap-frozen in liquid nitrogen. The pellet was suspended in 150 µl 2% Sodium Lauryl Sulfate (SLS) and proteins precipitated using acetone. For digestion, samples were resuspended in 0.5% SLS with 1 μg trypsin (Promega) and incubated for 30 min at 30°C, subsequently 5 mM TCEP were added to the suspension and further incubated overnight. Following, acetylation using 10 mM iodoacetamide for 30 min at 25°C in the dark, the peptides were desalted using C18 solid phase extraction. For label-free protein quantification, peptide mixtures were analysed using LC-MS. The data was acquired in data-independent acquisition mode and the MS raw data analysed by DIA-NN as described ^82, 83^. Data were further analysed and plotted using Python (3.7). The mass spectrometry proteomics data of whole cell proteomics and proximity labelling experiments have been deposited to the ProteomeXchange Consortium^84^ via the PRIDE^85^ partner repository with the dataset identifier PXD049046 (username; password: WxYzAbMr).

### Bioinformatics

The KEGG database ^86^ was used to assign functions to proteins, identify orthologs of *M. xanthus* proteins using a reciprocal best BlastP hit method and collect the 16s ribosomal RNA sequence of fully sequenced myxobacteria (Table S6). Protein domains were identified using InterPro ^87^, SMART ^88^, and the predicted AlphaFold structures. The DmxA protein sequence without the N-terminal transmembrane helices (amino acid 1-209) was used for AlphaFold-Multimer modelling via ColabFold (1.5.0) ^89–91^. The predicted Local Distance Difference Test (pLDDT) and predicted Alignment Error (pAE) graphs of the five models generated were made using a custom Matlab script. Models were ranked based on combined pLDDT and pAE values, with the best-ranked models used for further analysis and presentation. Per residue model accuracy was estimated based on pLDDT values (>90, high accuracy; 70-90, generally good accuracy; 50-70, low accuracy; <50, should not be interpreted) ^89^. Relative domain positions were validated by pAE. The pAE graphs indicate the expected position error at residue X if the predicted and true structures were aligned on residue Y; the lower the pAE value, the higher the accuracy of the relative position of residue pairs and, consequently, the relative position of domains/subunits/proteins ^89^. PyMOL version 2.4.1 (http://www.pymol.org/pymol) was used to analyse and visualize the models. The phylogenetic tree was prepared using the 16s ribosomal RNA sequence of fully sequenced myxobacteria in MEGA7 ^92^ using the Neighbor-Joining method ^93^. Bootstrap values (500 replicates) are shown next to the branches ^94^. RNA-seq. data were plotted using the BioMap function in Matlab. The base-by-base alignment coverage of RNA-seq and Cappable-seq reads of ^95^ were plotted for each position.

### Statistics

Colony expansion and c-di-GMP measurements were analysed using Student’s *t*-test in GraphPad Prism. Single cell speed and cell length distributions were analysed using the Mann-Whitney test in GraphPad Prism. Single cell reversal assays were analysed using the One-Way ANOVA function with Fishers LSD post-hoc test in GraphPad Prism. Asymmetry indexes (ω) were analysed using the rank-sum function (Mann-Whitney test) in Matlab. MAD was used as a measure of data variability and calculated based on the formula MAD=median(|x_i_ - x̃ |).

