## Supplementary Information for "A deterministic, c-di-GMP-dependent genetic program ensures the generation of phenotypically similar, symmetric daughter cells during cytokinesis"

**This file contains:**

- Supplementary Figures 1-6
- Supplementary Methods
- Table S1-S6
- Supplementary References

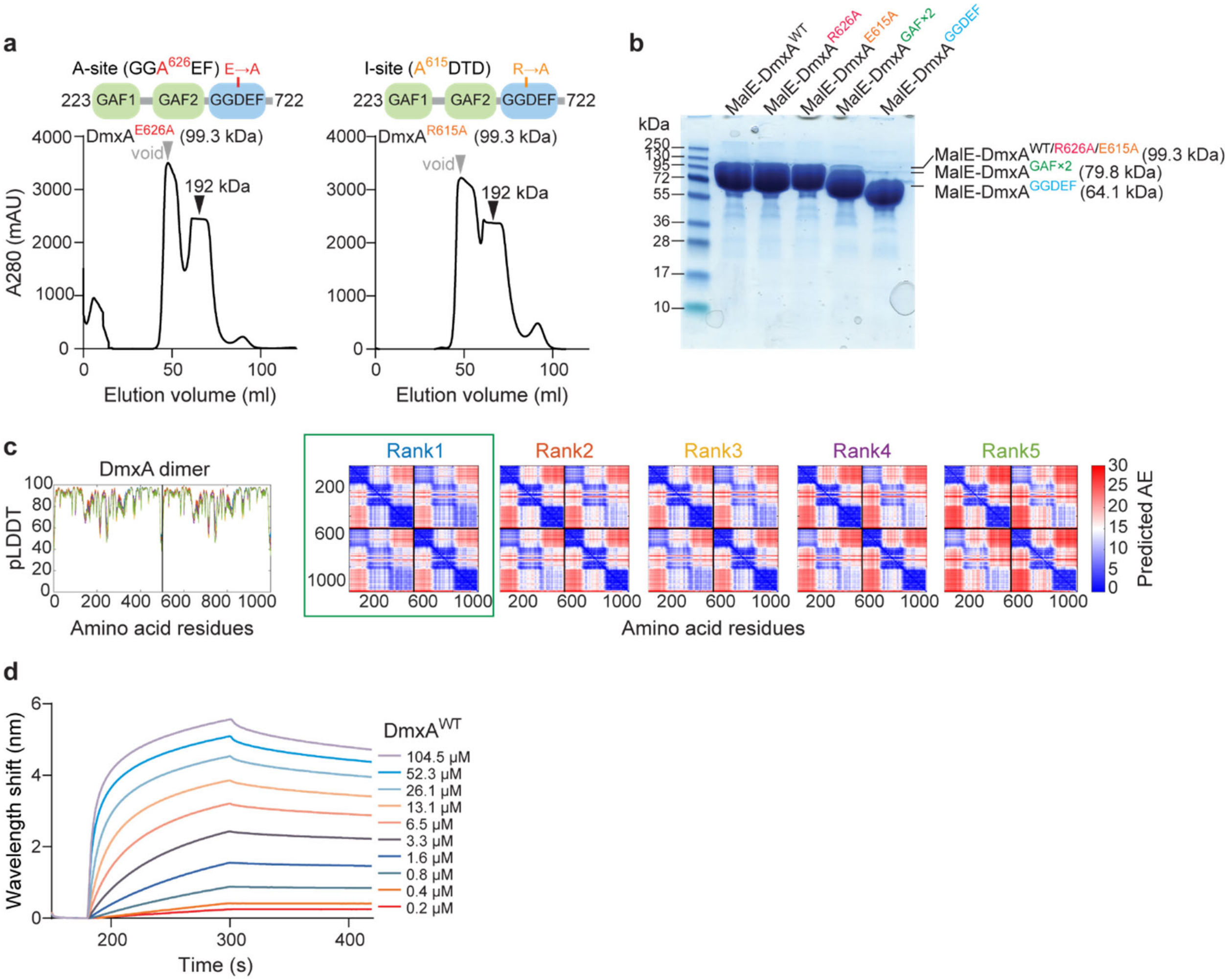

**Figure S1. *In silico* and *in vitro* analysis of DmxA variants.**

**a**, SEC of MalE-DmxA variants. Domain architectures of truncated DmxA variants are shown above chromatograms. Gray arrows indicate the void volume and black arrows the elution volume with the corresponding calculated molecular weight. **b**, Purification of MalE-tagged DmxA variants after SEC. Purified MalE-tagged DmxA variants were separated by SDS-PAGE and stained with Coomassie blue. Calculated molecular weights of proteins are indicated. Molecular size markers are indicated on the left. **c**, pLDDT and pAE plots for five models of the DmxA dimer structure in Fig. 1c predicted by AlphaFold-Multimer. The model marked by a green box was used for further analysis. **d**, Bio-Layer Interferometric analysis of the binding kinetics of MalE-tagged $DmxA^{WT}$ to c-di-GMP. Streptavidin-coated sensors were loaded with 500 nM biotinylated c-di-GMP and probed with indicated concentrations of $DmxA^{WT}$. The interaction kinetics were followed by monitoring the wavelength shifts during the association or dissociation of the analyte.

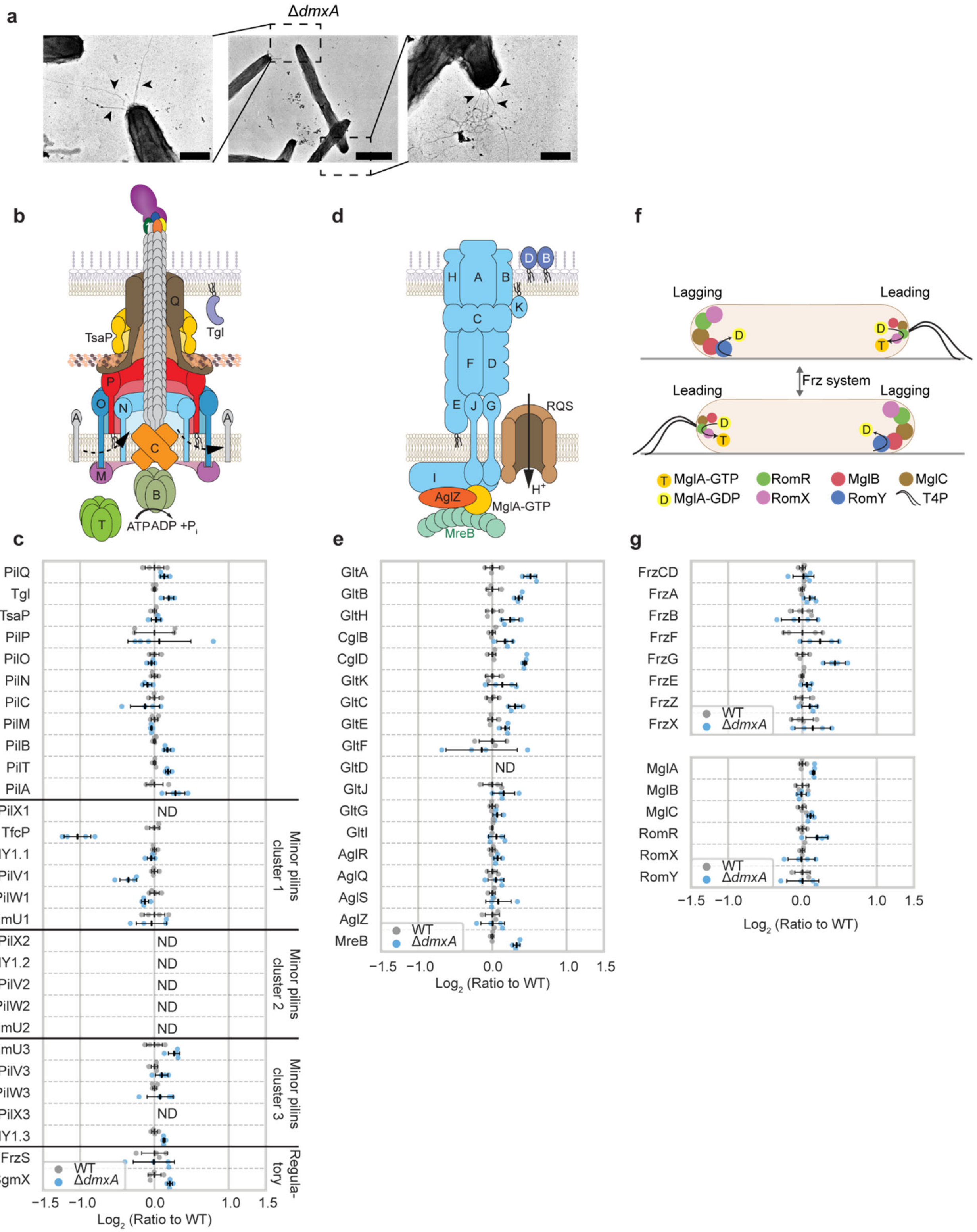

**Figure S2. Phenotypic characterization and proteome analysis of the Δ*dmxA* mutant.** **a**, Transmission electron microscopy of a Δ*dmxA* cell, with T4P at both poles. T4P are indicated by black arrowheads. The image in the middle shows the complete cell and the images on the left and right show the two poles. Scale bars, 0.5 µm (left and right), 2 µm (middle). **b, d, f**

Models of the architectural models of the T4PM (b) and gliding motility complex (d) as well as the localization of the proteins of the polarity module (f). In **b**, proteins labeled with single letters have the Pil prefix. The pilus fiber is formed by PilA subunits and a tip complex, composed of PilY1 and four minor pilins (blue: PilX, orange: PilV, green: PilW, yellow: FimU) [1, 2]. Tgl is an OM lipoprotein that is required for PilQ secretin assembly [3-5]. The ATPases PilB and PilT interact with the base of the T4PM in a mutually exclusive manner to stimulate extension and retraction of the pilus, respectively [1, 6, 7]. Bent arrows indicate incorporation of and removal of PilA subunits from the pilus base during extension and retraction, respectively. In **b** and **d**, lipoproteins are indicated by wavy black lines. In **d**, proteins labeled with single letters in light blue, dark blue or brown have the Glt, Cgl or Agl prefix, respectively [8-15]. The AglR/-Q/-S complex harnesses proton-motive force to power gliding motility [16, 17]. In **f**, proteins of the polarity module are indicated [18-26]. Upon signaling from the Frz chemosensory system, their polarity is inverted. For each protein, the size of a circle correlate with the amount of polarly localized protein. **c, e, g**, $Log_2$ fold-change of the accumulation of proteins of the T4PM (c), the Glt/Agl complex (e) for gliding, the Frz chemosensory system and the polarity module (g) in the Δ*dmxA* strain and WT cells compared to the mean of WT is shown. ND, not detected. Data points represent four biological replicates. Error bars, mean ± SD based on these four replicates.

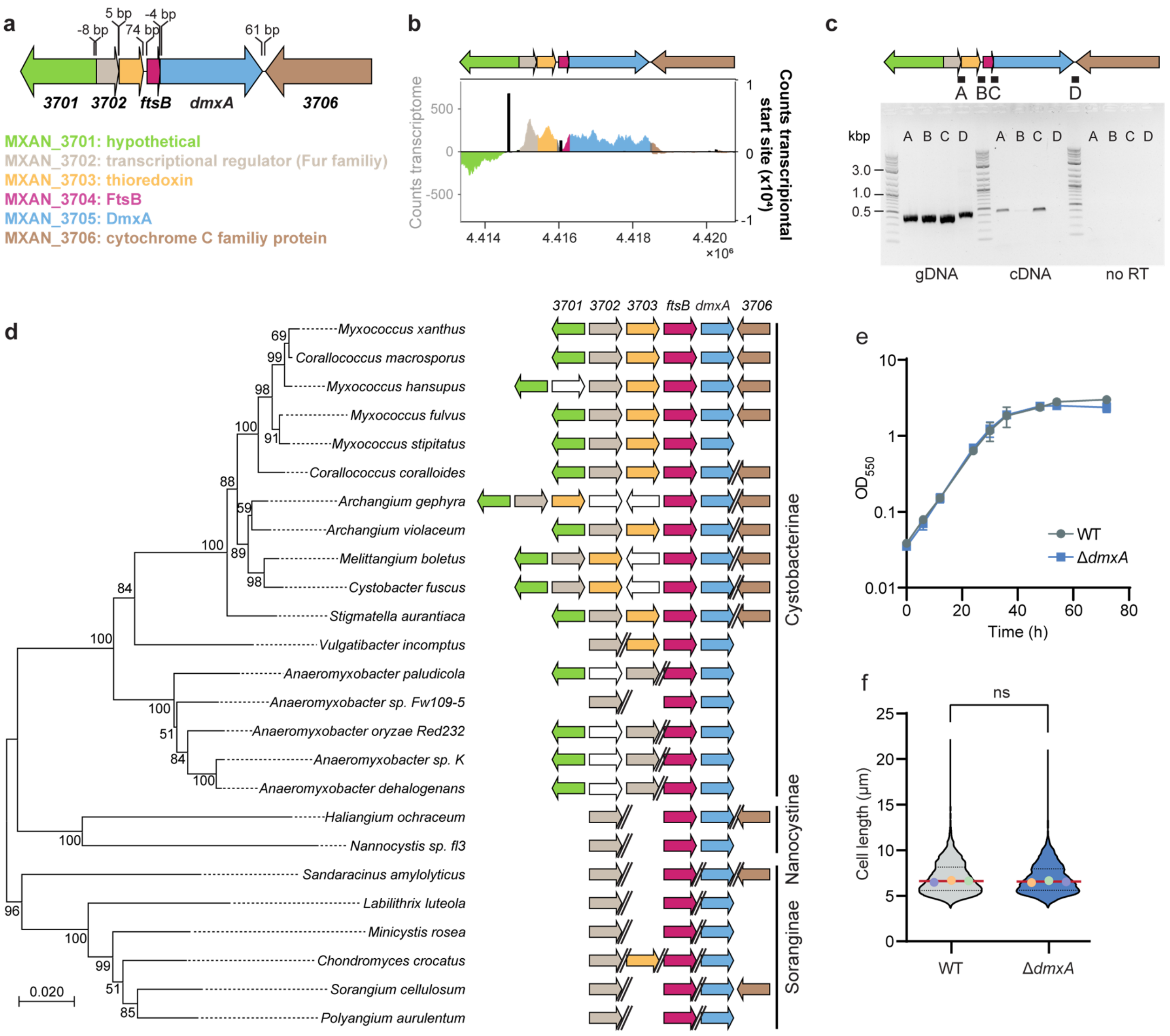

**Figure S3. Analysis of the *dmxA* locus.**

**a**, *dmxA* locus. Genes are drawn to scale and *MXAN* numbers or gene names are indicated. Predicted protein function is indicated. **b**, RNA-seq and Cappable-seq as base-by-base alignment coverage for total RNA isolated from cells growing in 1% CTT broth [27]. Upper panel, genetic organization of *dmxA* locus as in **a**. Lower panel, positive and negative values indicate reads mapped to the forward and reverse strand, respectively. Reads assigned to a gene are colored according to the gene colour code in **a**; intergenic regions are in gray. Cappable-seq coverage is shown as black bars. **c**, Operon mapping of the *dmxA* locus. Upper panel, genetic organization of *dmxA* locus as in **a**. Letters and black lines below the genes indicate the fragments amplified by PCR (~400-500 bp). Bottom panel, the PCR products amplified using genomic DNA, cDNA, and an enzyme-free reverse transcription reaction (no RT) as templates were separated on a 1% agarose gel. Letters above the individual lanes correspond to the letters of the primer combinations depicted above. Molecular size markers in kilo base-pairs (kbp) are shown on the left. **d**, Conservation of *dmxA* and its genetic neighborhood in other Myxococcales. Species included are listed in Table S6. Double slashes indicate no close proximity between the genes. **e**, Growth curve. Exponentially growing cells were diluted to an optical density (OD) at 550 nm ($OD_{550}$) of 0.04 and growth was followed over time. Growth curves were generated from three biological replicates. **f**, Cell length determination. The cell length distribution of three biological replicates is shown in a violin plot. Each violin indicates the probability density of the data at different cell length values. Single points represent the median of the three biological replicates (n=1652 in each of three biological replicates indicated in different colours). Median is represented by a continuous red line. Dashed lines indicate $25^{th}$ and $75^{th}$ percentiles. Samples were compared using a Mann-Whitney test; ns, not significant.

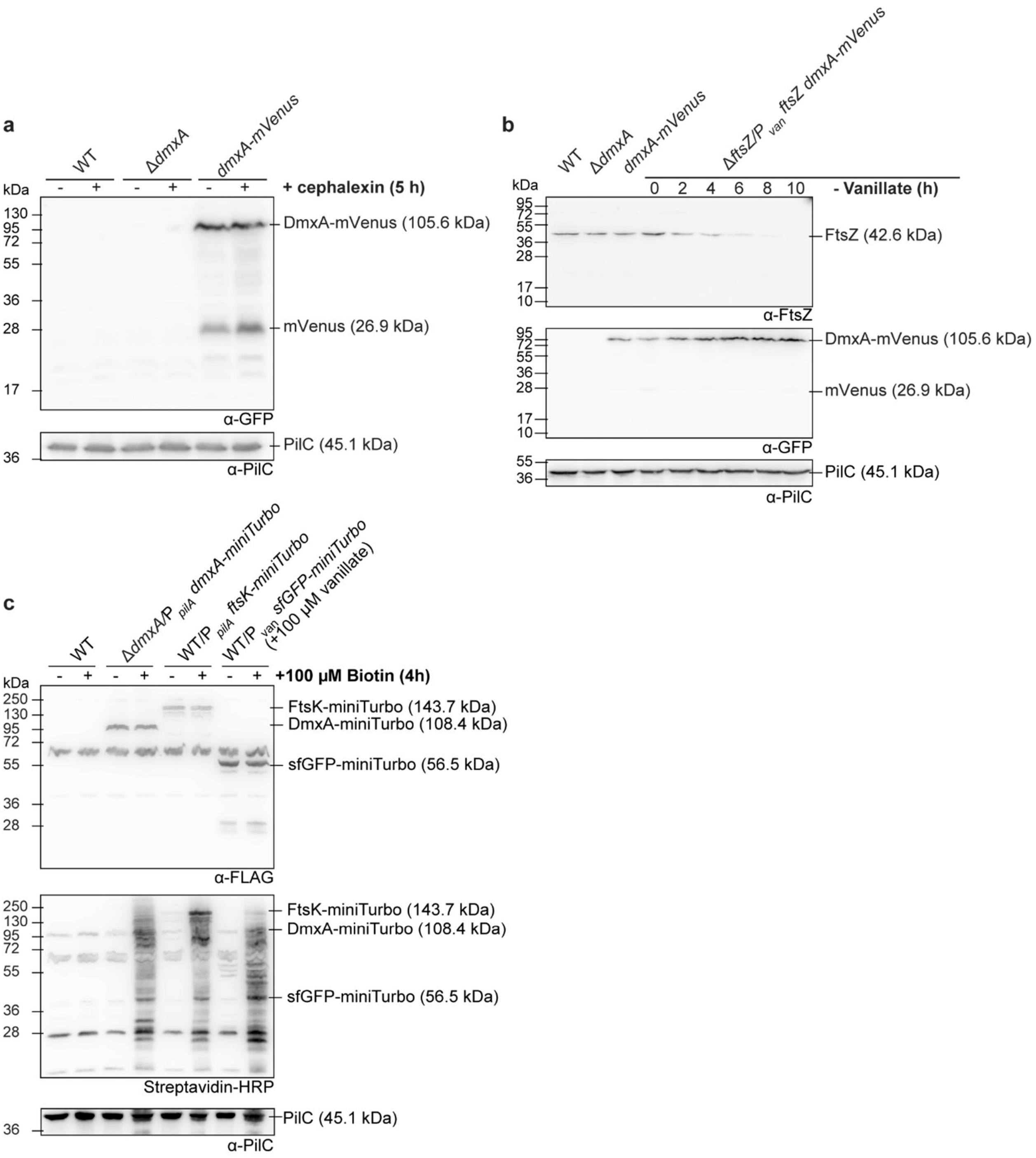

**Figure S4. Accumulation of DmxA-mVenus, FtsZ and miniTurbo variants.**
**a**, Immunoblot detection of DmxA-mVenus. Protein from 5.60×10$^{8}$ cells from exponentially growing suspension cultures treated with cephalexin for 5 h as indicated was loaded per lane. The same blot was stripped before applying a new antibody. PilC served as a loading control. Calculated molecular weights of proteins are indicated. **b**, Immunoblot detection of FtsZ and DmxA-mVenus. Protein from 1.05×10$^{8}$ cells from exponentially growing suspension cultures was loaded per lane. For the depletion of FtsZ, cells were exponentially grown in suspension culture in the presence of 10 μM vanillate for ~8 generations before the start of the experiment.

Then, cells were washed and subsequently grown for 10 h without vanillate. The same blot was stripped before applying a new antibody. PilC served as a loading control. Calculated molecular weights of proteins are indicated. **c**, Immunoblot detection of DmxA-, FtsK- and sfGFP-miniTurbo. DmxA-miniTurbo-FLAG and FtsK-miniTurbo-FLAG were expressed from the *pilA* promoter ($P_{pilA}$) and sfGFP-miniTurbo-FLAG from the $P_{van}$ in the presence of 100 µM vanillate. Protein from $7.00 \times 10^7$ cells from exponentially growing suspension cultures treated with or without biotin for 4 h was loaded per lane. The same blot was stripped before applying a new antibody. PilC served as a loading control.

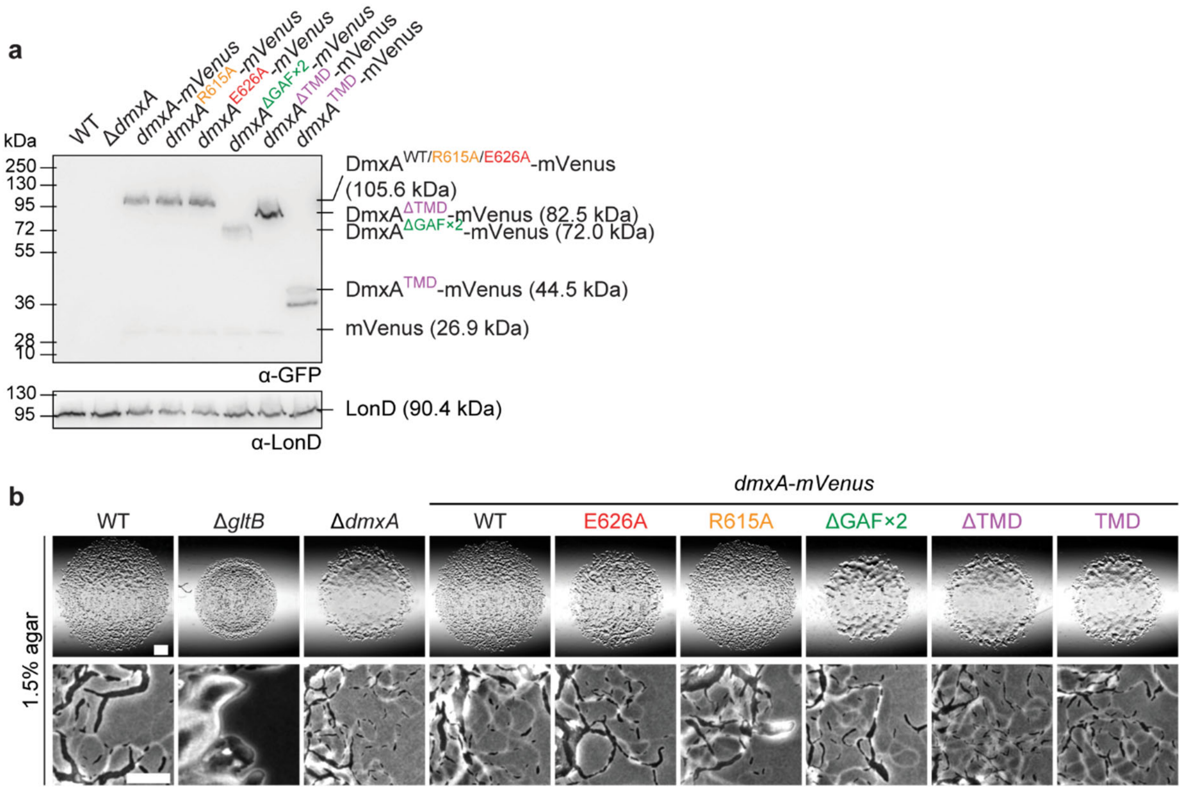

**Figure S5. Accumulation of DmxA-mVenus variants and gliding in their presence.**
**a**, Immunoblot detection of DmxA-mVenus variants. Protein from $1.40\times10^{8}$ cells from exponentially growing suspension cultures was loaded per lane. The same blot was stripped before applying a new antibody. LonD served as a loading control. Calculated molecular weights of proteins are indicated. **b**, Gliding was analyzed on 1.5% agar. Scale bars, 1 mm (upper panels), 50 µm (lower panels).

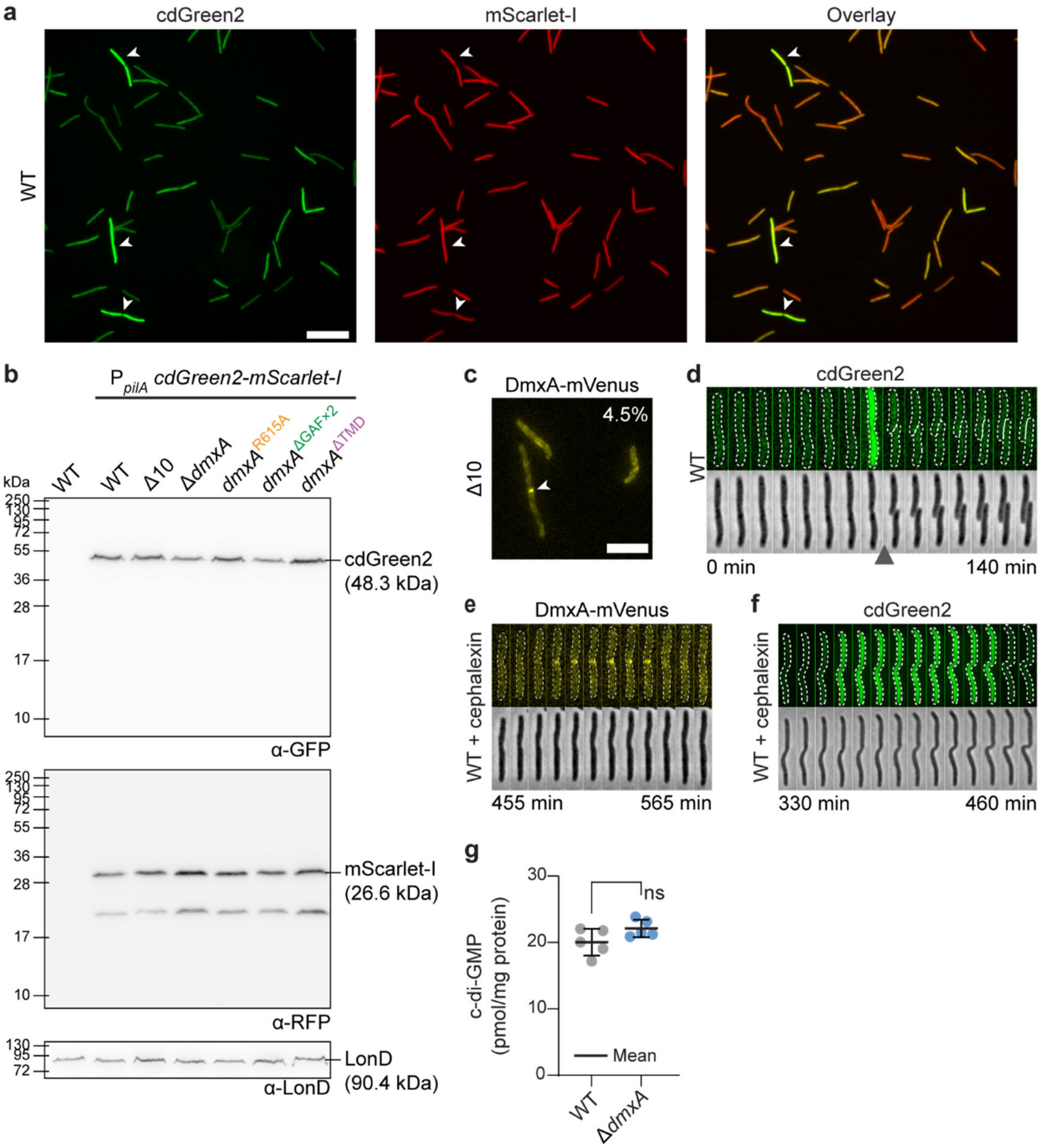

**Figure S6. Accumulation and analysis of cdGreen2 and mScarlet-I fluorescence**
**a**, Analysis of the cdGreen2 and mScarlet-I fluorescence in WT cells. White arrows indicate cells with bright cdGreen2 fluorescence. Scale bar, 10 µm. **b**, Immunoblot detection of cdGreen2 and mScarlet-I. Protein from $7.00\times10^{7}$ cells from exponentially growing suspension cultures was loaded per lane. The same blot was stripped before applying a new antibody. LonD served as a loading control. Calculated molecular weights of proteins are indicated. **c**, Localization of the DmxA-mVenus in Δ10 cells by epifluorescence microscopy. The percentage of cells with a cluster at mid-cell is indicated (n=200 from one biological replicate). White arrowhead indicate cluster. Scale bar, 5 µm. **d**, cdGreen2 fluorescence in WT cells during the cell cycle. Epifluorescence and phase-contrast images from time-lapse microscopy of cells expressing cdGreen2. Images were recorded every 10 min; arrowhead indicates completion of cytokinesis. **e-f**, Analysis of WT cells expressing DmxA-mVenus (e) or cdGreen2 (f) treated with

cephalexin. Images were recorded every 5 min (e) or every 10 min (f). **g**, c-di-GMP level of indicated strains during growth. The level of c-di-GMP is shown as the mean ± SD from five biological replicates. Individual data points are shown. ns, not significant, Student’s *t*-test.

**Supplementary Methods**

Plasmid construction. All oligonucleotides used are listed in Table S3. All constructed plasmids were verified by DNA sequencing.

**pMP072** (for in-frame deletion of *dmxA*): Up- and downstream fragments were amplified from genomic DNA using the primer pairs 3705_A/3705_B and 3705_C/3705_D, respectively. Subsequently, the up- and downstream fragments were used as a template for an overlapping PCR with the primer pair 3705_A/3705_D to generate the AD fragment. The AD fragment was digested with XbaI and KpnI, cloned in pBJ114 and sequenced. Of note, in order to successfully delete *dmxA*, the length of the flanking regions was increased compared to plasmid (pTP126) used in [28].

**pMP092** (plasmid for expression of *dmxA-mVenus* under control of $P_{nat}$ from the *attB* site): The $P_{nat}$ *dmxA* fragment was amplified from gDNA using the primer pair 3704 prmt forw +XbaI/3705_rev no stop 1. The mVenus fragment was amplified using pLC20 [22] as DNA template and the primer pair 3705_mVenus fw/mVenus_Kpn rev. To generate the full length insert, an overlapping PCR using the two fragments as DNA templates and the primer pair 3704 prmt forw +XbaI/mVenus_Kpn rev was performed. The fragment was digested with KpnI and XbaI, cloned into pSWU30 and sequenced.

**pMP093** (plasmid for replacement of *dmxA* with *dmxA-mVenus* in the native site): Up- and downstream fragments were amplified using genomic DNA from *M. xanthus* DK1622 as DNA template and the primer pairs 3705_native forw/3705_rev no stop 1 and 3705_native middle fw/3705_native rev, respectively. The mVenus fragment was amplified using pMP092 as DNA template and the primer pair 3705_mVenus fw/3705_native middle rev. To generate the full length insert, an overlapping PCR using the three fragments as DNA templates and the primer pair 3705_native forw/3705_native rev was performed. The fragment was digested with KpnI and XbaI, cloned into pBJ114 and sequenced.

**pMP164** (plasmid for expression of *dmxA-mVenus* from the native site in a Δ*dmxA* strain): Up- and downstream fragments were amplified using genomic DNA from *M. xanthus* DK1622 as a template and the primers 3705_A/3705_rev no stop 1 and 3705_native middle fw/3705_D respectively. mVenus was amplified from pMP093 as a template and the primers 3705_mVenus fw/3705_native middle rev. To generate the AD insert, an overlapping PCR using the three fragments as a DNA template and the primer pair 3705_A/3705_D was performed. The AD fragment was digested with KpnI and XbaI, cloned into pBJ114 and sequenced.

**pMP165** (plasmid for expression of *dmxA*$^{E626A}$*-mVenus* from the native site in a Δ*dmxA* strain): The mutation E626A was introduced into the plasmid pMP164 by using the primer pairs 3705_A/3705 E626A (-) and 3705 E626A (+)/3705_D and pMP164 as a DNA template for the PCR. To generate the AD insert, an overlapping PCR using both fragments as a DNA template and the primer pair 3705_A/3705_D was performed. The AD fragment was digested with KpnI and XbaI, cloned into pBJ114 and sequenced.

**pMP095** (plasmid for replacement of *dmxA* with *dmxA*$^{R615A}$*-mVenus* in the native site): Up- and downstream fragments were amplified using pMP093 as DNA template and the primer pairs 3705_native forw/3705 R615A (-) and 3705 R615A (+)/3705_native rev, respectively. To generate the full length insert, an overlapping PCR using the two fragments as DNA templates and the primer pair 3705_native forw/3705_native rev was performed. The fragment was digested with KpnI and XbaI, cloned into pBJ114 and sequenced.

**pKH02** (plasmid for deletion of *mVenus* in the *dmxA*$^{R615A}$*-mVenus* strain): Up- and downstream fragments were amplified using pMP095 as DNA template and the primer pairs 3705_native forw/3705_del_mVenus_(-) and 3705_del_mVenus_(+)/3705_native rev, respectively. To generate the full length insert, an overlapping PCR using the two fragments as DNA templates and the primer pair 3705_native forw/3705_native rev was performed. The fragment was digested with KpnI and XbaI, cloned into pBJ114 and sequenced.

**pMP175** (for generation of an in-frame deletion of the GAF domains of native *dmxA*): Up- and downstream fragments were amplified from pMP164 using the primer pairs 3705_PpilA forw/3705_B (GAFx1) and 3705_C (GAFx2)/3705_D (GAFx2), respectively. Subsequently, the up- and downstream fragments were used as a template for an overlapping PCR with the primer pair 3705_PpilA forw/3705_D (GAFx2) to generate the AD fragment. The AD fragment was digested with XbaI and KpnI, cloned in pBJ114 and sequenced.

**pMP179** (for generation of an in-frame deletion of the TMD domains of native *dmxA*): Up- and downstream fragments were amplified from pMP164 using the primer pairs 3705_A/3705_B3 (TMD) and 3705_C3 (TMD)/3705_D (TMD)2, respectively. Subsequently, the up- and downstream fragments were used as a template for an overlapping PCR with the primer pair 3705_A/3705_D (TMD)2 to generate the AD fragment. The AD fragment was digested with KpnI and XbaI, cloned in pBJ114 and sequenced.

**pMP182** (plasmid for expression of *dmxA*$^{TMD\ (1\text{-}167)}$-*mVenus* from the native site in a Δ*dmxA* strain): Up- and downstream fragments were amplified using pMP164 as a DNA template and the primers 3705_A/3705_TMH rev and 3705_mVenus fw2/3705_D respectively. To generate the AD insert, an overlapping PCR using the two fragments as a DNA template and the primer pair 3705_A/3705_D was performed. The AD fragment was digested with KpnI and XbaI, cloned into pBJ114 and sequenced.

**pMH113** (plasmid for expression of *malE-dmxA*$^{WT\ (223\text{-}722)}$): The *dmxA*$^{223\text{-}722}$ insert was amplified using genomic DNA from *M. xanthus* DK1622 as DNA template and the primer pair MalE-fwd_NotI/DmxA_Rev_HindIII. The fragment was digested with HindIII and NotI, cloned into pMAL-c6t and sequenced.

**pTP139** (plasmid for expression of His$_6$-*dmxA*$^{223\text{-}722,\ E626A}$): The *dmxA*$^{223\text{-}722,\ E626A}$ insert was amplified using pTP137 [28] as DNA template and the primer pairs 3705 GAF1 forw NdeI/3705 E626A (-) and 3705 E626A (+)/3705 rev BamHI. To generate the full length insert, an overlapping PCR using the two fragments as DNA templates and the primer pair 3705 GAF1 forw NdeI/3705 rev BamHI was performed. The fragment was digested with NdeI and BamHI, cloned into pET28a(+) and sequenced.

**pMH117** (plasmid for expression of *malE-dmxA*$^{E626A\ (223\text{-}722,\ E626A)}$): The *dmxA*$^{223\text{-}722,\ E626A}$ insert was amplified using pTP139 as DNA template and the primer pair MalE-fwd_NotI/DmxA_Rev_HindIII. The fragment was digested with HindIII and NotI, cloned into pMAL-c6t and sequenced.

**pMP082** (plasmid for expression of His$_6$-*dmxA*$^{223\text{-}722,\ R615A}$): The *dmxA*$^{223\text{-}722,\ R615A}$ insert was amplified using pTP137 [28] as DNA template and the primer pairs 3705 GAF1 forw NdeI/3705 R615A (-) and 3705 R615A (+)/3705 rev BamHI. To generate the full length insert, an overlapping PCR using the two fragments as DNA templates and the primer pair 3705 GAF1 forw NdeI/3705 rev BamHI was performed. The fragment was digested with NdeI and BamHI, cloned into pET28a(+) and sequenced.

**pMH116** (plasmid for expression of *malE-dmxA*$^{R615A\ (223\text{-}722,\ R615A)}$): The *dmxA*$^{223\text{-}722,\ R615A}$ insert was amplified using pMP082 as DNA template and the primer pair MalE-fwd_NotI/DmxA_Rev_HindIII. The fragment was digested with HindIII and NotI, cloned into pMAL-c6t and sequenced.

**pMH114** (plasmid for expression of *malE-dmxA*$^{GAF\times2\ (223\text{-}547)}$): The *dmxA*$^{223\text{-}547}$ insert was amplified using genomic DNA from *M. xanthus* DK1622 as DNA template and the primer pair MalE-fwd_NotI/DmxA_GAF2_rev_HindIII. The fragment was digested with HindIII and NotI, cloned into pMAL-c6t and sequenced.
**pMH115** (plasmid for expression of *malE-dmxA*$^{GGDEF\ (546\text{-}722)}$): The *dmxA*$^{546\text{-}722}$ insert was amplified using genomic DNA from *M. xanthus* DK1622 as DNA template and the primer pair MalE-fwd_NotI/DmxA_GGDEFonly_Fwd_NotI. The fragment was digested with HindIII and NotI, cloned into pMAL-c6t and sequenced.
**pMAT74** (for in-frame deletion of *ftsZ*): up- and downstream fragments were amplified using genomic DNA from *M. xanthus* DK1622 as DNA template and the primer pairs ftsZ-up EcoRI/ftsZ-overlapping reverse and ftsZ-overlapping forward/ftsZ-down HindIII, respectively. Subsequently, the up- and downstream fragments were used as a template for an overlapping PCR with the primer pair ftsZ-up EcoRI/ftsZ-down HindIII to generate the AD fragment. The AD fragment was digested with EcoRI and HindIII, cloned in pBJ114 and sequenced.
**pMAT86** (plasmid for expression of *ftsZ* from the *MXAN_18-19* site under the control of the vanillate promoter): the *ftsZ* fragment was amplified using genomic DNA from *M. xanthus* DK1622 as DNA template and the primer pair ftsZ-start NdeI/ftsZ-stop KpnI. The fragment was digested with NdeI and KpnI, cloned into pMR3691 and sequenced.
**pMH52** (plasmid containing a *M. xanthus* codon optimized *miniTurbo-FLAG*) The *miniTurbo* sequence [29] was codon optimized for *M. xanthus*, synthesized with a N-terminal GGGS-linker and an C-terminal FLAG-tag and cloned into the pEX-k168.
**pMP172** (plasmid for expression of *dmxA-miniTurbo-FLAG* under control of the *pilA* promoter from the *attB* site): The *dmxA* fragment was amplified with the primer pair 3705_PpilA forw/3705_rev no stop 1 from pMP092, and the *miniTurbo-FLAG* fragment was amplified with the primer pair 3705_miniTurboID fw/Flag_rev HindIII from pMH52. Next, and overlapping PCR was performed using the previous PCR products and the primer pair 3705_PpilA forw/Flag_rev HindIII. The product was digested with XbaI and HindIII, cloned into pSW105 and sequenced.
**pMH98** (plasmid for replacement of *ftsK* with *ftsK-miniTurbo-FLAG* in the native site): up- and downstream fragments were amplified using genomic DNA from *M. xanthus* DK1622 as DNA template and the primers FtsK_A_BamHI/FtsK_B_TID_OV and FtsK_C_TID_OV/FtsK_D_HindIII, respectively. The *miniTurbo-FLAG* fragment was amplified using pMH52 and the primer pair TID_fwd_FtsK_OV/TID_rev_FtsK_OV. To generate the full length insert, an overlapping PCR using the three fragments as DNA templates and the primer pair FtsK_A_BamHI/FtsK_D_HindIII was performed. The fragment was digested with BamHI and HindIII, cloned into pBJ114 and sequenced.
**pMH110** (plasmid for expression of *ftsK-miniTurbo-FLAG* from the *attB* site under the control of the *pilA* promoter): *ftsK* was amplified using genomic DNA from *M. xanthus* DK1622 as DNA template and the primer pair FtsK_fwd_XbaI_2/FtsK_B_TID_OV 2. The *miniTurbo-FLAG* insert was amplified from pMH98 using the primer pair TID_fwd_FtsK_OV/Flag_rev HindIII. To generate the full length insert, an overlapping PCR using the two fragments as DNA templates and the primer pair FtsK_fwd_XbaI_2/Flag_rev HindIII was performed. The fragment was digested with XbaI and HindIII, cloned into pSW105 and sequenced.
**pMH123** (plasmid for expression of codon optimized cdGreen2 biosensor and mScarlet-I from the *attB* site under the control of the *pilA* promoter): The codon-optimized cdGreen2 biosensor and mScarlet-I full-length insert was amplified using pBBR15.2-2H12.D11opt-scarRef [30] as DNA

template and the primer pair cdG-Sensor_fwd_XbaI/cdG-Sensor_rev_HindIII. The fragment was digested with XbaI and HindIII, cloned into pSW105 and sequenced.
**pMEM33** (plasmid for replacement of *pilT* with *mCherry-pilT* in the native site): the fragment was amplified using pAP87 [31] as a DNA template and the primer primer pair PilT fw EcoRI/PilT rev HindIII. The fragment was digested with EcoRI and HindIII, cloned into pBJ114 and sequenced.
**pMEM23** (plasmid for replacement of *pilB* with *pilB-mCherry* in the native site): the *pilB-mCherry* fragment was amplified using pAP12 [31] as a DNA template and the primer primer pairs pilB-mCA-EcoRI/pilB-mc-B-overlay. The downstream fragment of *pilB* was amplified using genomic DNA from *M. xanthus* DK1622 and the primer pair pilB-C-overlay/pilB-D-HindIII. To generate the full length insert, an overlapping PCR using the two fragments as DNA templates and the primer pair pilB-mCA-EcoRI/pilB-D-HindIII was performed. The fragment was digested with EcoRI and HindIII, cloned into pBJ114 and sequenced.

**Table S1.** Proteins significantly enriched in DmxA-miniTurbo-FLAG proximity labeling experiments

| Locus Tag | Name | Annotation | log2Ratio* | -log10(pValue)** |
|---|---|---|---|---|
| MXAN_3705 | DmxA | Diguanylate cyclase | 15.15 | 6.13 |
| MXAN_5591 | | FHA domain/tetratricopeptide repeat protein | 6.20 | 5.63 |
| MXAN_4591 | Pkn1 | Serine/threonine-protein kinase | 5.87 | 5.81 |
| MXAN_5723 | | MJ0042 family finger-like domain protein | 5.36 | 3.56 |
| MXAN_0755 | Pkn9 | Serine/threonine protein kinase | 5.28 | 3.25 |
| MXAN_3955 | Pkn10 | Serine/threonine protein kinase | 5.16 | 5.40 |
| MXAN_4324 | | Uncharacterized protein | 5.03 | 2.13 |
| MXAN_1436 | | Hydrolase, alpha/beta fold family | 4.95 | 4.13 |
| MXAN_1131 | | Uncharacterized protein | 4.74 | 3.69 |
| MXAN_4024 | | Uncharacterized protein | 4.64 | 3.48 |
| MXAN_6266 | | Putative 2,3-cyclic-nucleotide 2-phosphodiesterase | 4.58 | 2.74 |
| MXAN_4956 | | Uncharacterized protein | 4.45 | 5.56 |
| MXAN_2680 | | Serine/threonine protein kinase | 4.43 | 4.74 |
| MXAN_1350 | | Histidine kinase | 4.34 | 3.88 |
| MXAN_5778 | PilS2 | Histidine kinase | 4.32 | 4.86 |
| MXAN_5946 | | Cytochrome c family protein/iron permease, FTR1 family | 4.23 | 2.86 |
| MXAN_4677 | | Uncharacterized protein | 4.21 | 2.98 |
| MXAN_3550 | | Uncharacterized protein | 4.18 | 3.96 |
| MXAN_7417 | EpsY | OPX polysaccharide export protein | 4.18 | 2.05 |
| MXAN_4735 | | Putative membrane protein | 4.16 | 3.20 |
| MXAN_6605 | CsdK1 | PilZ-domain/DnaK family protein | 4.16 | 4.27 |
| MXAN_2059 | | Putative serine/threonine protein kinase | 4.14 | 4.39 |
| MXAN_1134 | | Adenylate/guanylate cyclase domain protein | 4.12 | 3.31 |
| MXAN_0117 | | Serine/threonine protein kinase | 4.05 | 2.13 |
| MXAN_4700 | | Serine/threonine protein kinase | 4.00 | 3.67 |
| MXAN_1622 | | Uncharacterized protein | 3.98 | 3.40 |
| MXAN_5710 | | Pyrimidine-nucleoside phosphorylase | 3.86 | 2.38 |
| MXAN_3108 | | FHA domain/tetratricopeptide repeat protein | 3.79 | 2.47 |
| MXAN_5321 | | Uncharacterized protein | 3.65 | 4.04 |
| MXAN_3106 | | Protein transporter, outer bacterial membrane secretin (Secretin) family | 3.64 | 4.98 |
| MXAN_2710 | | Putative lipoprotein | 3.59 | 4.85 |
| MXAN_1460 | FtsK | Cell division protein | 3.56 | 2.62 |
| MXAN_1942 | | MJ0042 family finger-like domain/tetratricopeptide repeat protein | 3.46 | 4.37 |
| MXAN_6420 | Pkn2 | Serine/threonine protein kinase | 3.45 | 2.68 |
| MXAN_5667 | | Hydrolase, alpha/beta fold family | 3.44 | 3.96 |
| MXAN_2520 | | FHA domain/tetratricopeptide repeat protein | 3.39 | 4.06 |
| MXAN_3343 | | Histidine kinase | 3.32 | 3.44 |
| MXAN_1249 | | Histidine kinase | 3.30 | 3.03 |

| | | | | |
|---|---|---|---|---|
| MXAN_3778 | CsdK2 | PilZ-domain/DnaK family protein | 3.27 | 3.17 |
| MXAN_4730 | | Lipoprotein releasing system, transmembrane protein, LolC/E family | 3.25 | 2.87 |
| MXAN_2683 | | Methyl accepting chemotaxis protein | 3.23 | 3.02 |
| MXAN_5813 | | Uncharacterized protein | 3.22 | 4.45 |
| MXAN_6183 | | Serine/threonine protein kinase | 3.14 | 2.97 |
| MXAN_3808 | | Protease 4 | 3.14 | 2.49 |
| MXAN_3159 | | Uncharacterized protein | 3.14 | 4.50 |
| MXAN_2504 | | Putative general secretion pathway protein L | 3.12 | 4.42 |
| MXAN_3461 | | Oxidoreductase, short chain dehydrogenase/reductase family | 3.11 | 2.20 |
| MXAN_6720 | | Putative lipoprotein | 3.08 | 4.70 |
| MXAN_1313 | | Uncharacterized protein | 3.04 | 3.11 |
| MXAN_0745 | | Uncharacterized protein | 3.03 | 4.34 |

[*] Fold change between the average intensities of the DmxA-miniTurbo-FLAG and the sfGFP-miniTurbo-FLAG samples.

[**] $Log_{10}$ of p-value.

**Table S2.** Proteins significantly enriched in FtsK-miniTurbo-FLAG proximity labeling experiments

| Locus Tag | Name | Annotation | log2Ratio* | -log10(pValue)** |
|---|---|---|---|---|
| MXAN_1460 | FtsK | Cell division protein | 10.85 | 6.07 |
| MXAN_5723 | | MJ0042 family finger-like domain protein | 6.34 | 3.87 |
| MXAN_5591 | | FHA domain/tetratricopeptide repeat protein | 6.17 | 6.27 |
| MXAN_2297 | | Uncharacterized protein | 5.95 | 3.90 |
| MXAN_0960 | | Putative serine/threonine protein kinase | 5.82 | 2.07 |
| MXAN_4359 | FtsH | ATP-dependent zinc metalloprotease | 5.79 | 3.96 |
| MXAN_3108 | | FHA domain/tetratricopeptide repeat protein | 5.66 | 4.81 |
| MXAN_4700 | | Serine/threonine protein kinase | 5.65 | 4.83 |
| MXAN_2438 | | Type III secretion apparatus protein, YscI/HrpB family | 5.34 | 2.94 |
| MXAN_2710 | | Putative lipoprotein | 5.34 | 5.51 |
| MXAN_3705 | DmxA | Diguanylate cyclase | 5.25 | 3.34 |
| MXAN_1527 | | NAD dependent epimerase/dehydratase family protein | 5.22 | 5.24 |
| MXAN_2077 | | Protein kinase domain protein | 5.07 | 3.79 |
| MXAN_4037 | | Cytidylate kinase | 4.98 | 3.80 |
| MXAN_3106 | | Protein transporter, outer bacterial membrane secretin (Secretin) family | 4.90 | 5.09 |
| MXAN_0363 | TfcP | T4P motility protein | 4.80 | 5.48 |
| MXAN_6605 | CsdK1 | PilZ-domain/DnaK family protein | 4.78 | 4.31 |
| MXAN_6860 | AglS | Gliding motility protein | 4.78 | 2.82 |
| MXAN_1419 | | Uncharacterized protein | 4.77 | 3.92 |
| MXAN_4186 | | Uncharacterized protein | 4.71 | 3.46 |
| MXAN_5765 | | Putative general secretion pathway protein A | 4.69 | 4.47 |
| MXAN_6224 | | DNA-binding response regulator, Fis family | 4.68 | 4.14 |
| MXAN_5772 | PilQ | T4P secretin | 4.66 | 3.77 |
| MXAN_5028 | | Putative membrane protein | 4.59 | 4.64 |
| MXAN_3001 | TsaP | T4P motility protein | 4.58 | 6.12 |
| MXAN_0755 | Pkn9 | Serine/threonine protein kinase | 4.57 | 2.84 |
| MXAN_4024 | | Uncharacterized protein | 4.51 | 3.17 |
| MXAN_5075 | | Ribose-phosphate pyrophosphokinase | 4.49 | 3.01 |
| MXAN_5319 | PglC | Tetratricopeptide repeat protein | 4.43 | 5.96 |
| MXAN_6705 | PilT5 | T4P retraction ATPase | 4.41 | 4.04 |
| MXAN_3013 | | ATP-dependent protease ATPase subunit HslU | 4.37 | 5.57 |
| MXAN_4667 | | General secretion pathway protein E, N-terminal domain protein | 4.32 | 4.10 |
| MXAN_6183 | | Serine/threonine protein kinase | 4.29 | 3.94 |
| MXAN_6861 | AglQ | Gliding motility protein | 4.26 | 3.75 |
| MXAN_4737 | | HDIG domain protein | 4.17 | 3.51 |
| MXAN_2968 | | Efflux transporter, RND family, MFP subunit | 4.12 | 5.30 |

| | | | | |
|---|---|---|---|---|
| MXAN_6420 | Pkn2 | Serine/threonine protein kinase | 4.11 | 4.07 |
| MXAN_3202 | Pkn3 | Serine/threonine protein kinase | 4.05 | 5.55 |
| MXAN_1361 | | FHA domain protein | 4.04 | 2.87 |
| MXAN_2312 | | Uncharacterized protein | 4.04 | 3.42 |
| MXAN_1350 | | Histidine kinase | 4.02 | 5.82 |
| MXAN_0459 | | Histidine kinase | 3.98 | 4.29 |
| MXAN_3337 | | NAD-dependent protein deacylase | 3.95 | 3.38 |
| MXAN_5597 | FtsZ | Cell division protein | 3.93 | 3.86 |
| MXAN_3550 | | Uncharacterized protein | 3.91 | 5.69 |
| MXAN_2483 | | Type II DNA topoisomerase, B subunit | 3.91 | 4.74 |
| MXAN_7208 | | Serine/threonine protein kinase | 3.90 | 4.99 |
| MXAN_2480 | | Signal peptide peptidase SppA, 36K type | 3.89 | 3.75 |
| MXAN_2550 | Pkn6 | Serine/threonine protein kinase | 3.85 | 2.98 |
| MXAN_3710 | | Serine/threonine protein kinase | 3.83 | 2.02 |
| MXAN_5152 | | OmpA family protein | 3.83 | 5.40 |
| MXAN_5735 | FtsY | Signal recognition particle receptor | 3.79 | 5.32 |
| MXAN_2680 | | Serine/threonine protein kinase | 3.79 | 4.07 |
| MXAN_1178 | | FKBP-type peptidyl-prolyl isomerase, trigger factor family | 3.78 | 2.91 |
| MXAN_5783 | PilA | T4P major pilin | 3.77 | 5.24 |
| MXAN_7025 | | DnaK family protein | 3.76 | 3.88 |
| MXAN_4371 | | Serine/threonine protein kinase | 3.73 | 3.44 |
| MXAN_2586 | | Serine/threonine protein kinase | 3.72 | 5.00 |
| MXAN_0625 | | Aldehyde dehydrogenase | 3.69 | 2.37 |
| MXAN_1948 | | TPR domain protein | 3.65 | 5.25 |
| MXAN_5786 | PilC | T4P motility protein | 3.57 | 3.67 |
| MXAN_7371 | | Serine/threonine protein kinase | 3.54 | 3.43 |
| MXAN_5129 | | Uncharacterized protein | 3.52 | 4.58 |
| MXAN_4094 | | Uncharacterized protein | 3.48 | 3.27 |
| MXAN_0018 | | Serine/threonine protein kinase | 3.48 | 3.78 |
| MXAN_6704 | | Acetyltransferase, GNAT family | 3.40 | 3.37 |
| MXAN_0882 | | Serine/threonine protein kinase | 3.37 | 3.68 |
| MXAN_3570 | | Uncharacterized protein | 3.37 | 4.62 |
| MXAN_4735 | | Putative membrane protein | 3.36 | 4.36 |
| MXAN_1984 | | Uncharacterized protein | 3.34 | 4.76 |
| MXAN_0653 | | Peptidase, S8A (Subtilisin) subfamily | 3.33 | 3.38 |
| MXAN_6310 | | AhpC/TSA family protein | 3.33 | 2.08 |
| MXAN_2444 | | Uncharacterized protein | 3.27 | 2.86 |
| MXAN_1995 | PilT | T4P retraction ATPase | 3.27 | 3.01 |
| MXAN_2512 | GspF | General secretion pathway protein F | 3.25 | 3.15 |
| MXAN_6570 | | Serine/threonine protein kinase | 3.22 | 2.82 |
| MXAN_0344 | | Metallophosphoesterase | 3.16 | 3.89 |

| | | | | |
|---|---|---|---|---|
| MXAN_6155 | | Alkylphosphonate utilization operon protein PhnA | 3.13 | 5.22 |
| MXAN_0871 | | Putative serine/threonine protein kinase | 3.13 | 2.30 |
| MXAN_5696 | | Serine/threonine protein kinase | 3.13 | 3.87 |
| MXAN_4591 | Pkn1 | Serine/threonine protein kinase | 3.12 | 3.72 |
| MXAN_6189 | | Uncharacterized protein | 3.12 | 3.63 |
| MXAN_4867 | GltF | Gliding motility protein | 3.12 | 4.51 |
| MXAN_6611 | | Uncharacterized protein | 3.11 | 3.34 |
| MXAN_2013 | | Trigger factor | 3.06 | 5.01 |
| MXAN_0599 | | Uncharacterized protein | 3.06 | 2.33 |
| MXAN_5824 | | Putative membrane protein | 3.06 | 3.67 |
| MXAN_3984 | | SPFH/band 7 domain protein | 3.04 | 2.40 |
| MXAN_3918 | | Glutamate synthase, small subunit | 3.04 | 3.80 |
| MXAN_6751 | | Uncharacterized protein | 3.03 | 4.12 |
| MXAN_2156 | | Serine/threonine protein kinase | 3.03 | 3.98 |

[*] Fold change between the average intensities of the FtsK-miniTurbo-FLAG and the sfGFP-miniTurbo-FLAG samples.

[**] $Log_{10}$ of p-value.

**Table S3.** Strains used in this work

| Strain | Genotype | Reference |
|---|---|---|
| ***M. xanthus*** | | |
| DK1622 | Wildtype | [32] |
| DK10410 | Δ*pilA* | [33] |
| SA3922 | Δ*gltB* | [8] |
| SA8802 | Δ*frzE* | [22] |
| SA3387 | Δ*mglB* | [20] |
| SA7442 | Δ*dmxA* | This study |
| SA7447 | Δ*dmxA attB::*pTP140 ($P_{nat}$ *dmxA*) | This study |
| SA7466 | Δ*dmxA* Δ*pilA* | This study |
| SA12010 | Δ*dmxA* Δ*gltB* | This study |
| SA7479 | Δ*dmxA* Δ*frzE* | This study |
| SA7485 | *dmxA-mVenus* | This study |
| SA8589 | *dmxA-mVenus* Δ*gltB* | This study |
| SA8580 | *dmxA*$^{E626A}$*-mVenus* | This study |
| SA7488 | *dmxA*$^{R615A}$*-mVenus* | This study |
| SA12015 | *dmxA*$^{\Delta GAF\times 2\ (\Delta 223\text{-}532)}$*-mVenus* | This study |
| SA12034 | *dmxA*$^{\Delta TMD\ (\Delta 3\text{-}222)}$*-mVenus* | This study |
| SA12065 | *dmxA*$^{TMD\ (1\text{-}167)}$*-mVenus* | This study |
| SA6755 | Δ*ftsZ MXAN_18-19::*pMAT86 ($P_{van}$ *ftsZ*) | This study |
| SA8511 | Δ*ftsZ MXAN_18-19::*pMAT86 ($P_{van}$ *ftsZ*) *dmxA-mVenus* | This study |
| SA8596 | Δ*dmxA attB::*pMP172 ($P_{pilA}$ *dmxA-miniTurbo-FLAG*) | This study |
| SA12038 | *attB::*pMH110 ($P_{pilA}$ *ftsK-miniTurbo-FLAG*) | This study |
| SA12027 | WT *MXAN_18-19::*pMH97 ($P_{van}$ *sfGFP-miniTurbo-FLAG*) | [34] |
| SA9082 | *attB::*pMH123 ($P_{pilA}$ *cdGreen2 mScarlet-I*) | This study |
| SA12093 | Δ*gltB attB::*pMH123 ($P_{pilA}$ *cdGreen2 mScarlet-I*) | This study |
| SA12091 | Δ*dmxA attB::*pMH123 ($P_{pilA}$ *cdGreen2 mScarlet-I*) | This study |
| SA12092 | Δ*dmxA* Δ*gltB attB::*pMH123 ($P_{pilA}$ *cdGreen2 mScarlet-I*) | This study |
| SA5666 | Δ10 (Δ*MXAN_1525*, Δ*MXAN_4029*, Δ*MXAN_5199*, Δ*MXAN_7362,* Δ*gacB* (Δ*MXAN_4463*), Δ*gacA* (Δ*MXAN_2643*), Δ*MXAN_2997*, Δ*MXAN_5366*, Δ*dmxB* (Δ*MXAN_3735*), Δ*MXAN_5791*) | This study |
| SA12098 | Δ10* *dmxA-mVenus* | This study |
| SA9083 | Δ10* *attB::*pMH123 ($P_{pilA}$ *cdGreen2 mScarlet-I*) | This study |
| SA12063 | Δ10* Δ*gltB* | This study |
| SA9084 | Δ10* Δ*gltB attB::*pMH123 ($P_{pilA}$ *cdGreen2 mScarlet-I*) | This study |
| SA13309 | *dmxA*$^{R615A}$ | This study |
| SA13310 | *dmxA*$^{R615A}$ *attB::*pMH123 ($P_{pilA}$ *cdGreen2 mScarlet-I*) | This study |
| SA12023 | *dmxA*$^{\Delta GAF\times 2\ (\Delta 223\text{-}532)}$ | This study |
| SA12097 | *dmxA*$^{\Delta GAF\times 2\ (\Delta 223\text{-}532)}$ *attB::*pMH123 ($P_{pilA}$ *cdGreen2 mScarlet-I*) | This study |
| SA12029 | *dmxA*$^{\Delta TMD\ (\Delta 3\text{-}222)}$ | This study |
| SA12095 | *dmxA*$^{\Delta TMD\ (\Delta 3\text{-}222)}$ *attB::*pMH123 ($P_{pilA}$ *cdGreen2 mScarlet-I*) | This study |
| SA7192 | *pilQ-sfGFP* | [31] |
| SA12017 | *pilQ-sfGFP* Δ*gltB* | [5] |

| | | |
|---|---|---|
| SA8505 | Δ*dmxA pilQ-sfGFP* | This study |
| SA12019 | Δ*dmxA pilQ-sfGFP* Δ*gltB* | This study |
| SA7896 | *mCherry-pilM* | [2] |
| SA8576 | Δ*dmxA mCherry-pilM* | This study |
| SA9307 | *mCherry-pilT* | This study |
| SA8575 | Δ*dmxA mCherry-pilT* | This study |
| SA9300 | *pilB-mCherry* | This study |
| SA8572 | Δ*dmxA pilB-mCherry* | This study |
| SA7507 | *romR-mCherry* | [22] |
| SA12056 | *romR-mCherry* Δ*gltB* | This study |
| SA7459 | Δ*dmxA romR-mCherry* | This study |
| SA12064 | Δ*dmxA romR-mCherry* Δ*gltB* | This study |
| SA10043 | *mglB-mVenus* | [23] |
| SA12024 | Δ*dmxA mglB-mVenus* | This study |
| SA8185 | *mglA-mVenus* | [22] |
| SA7480 | Δ*dmxA mglA-mVenus* | This study |
| SA7195 | *sgmX-mVenus* | [31] |
| SA12014 | Δ*dmxA sgmX-mVenus* | This study |
| ***E. coli*** | | |
| Mach1 | Δ*recA*1398 *endA*1 *tonA* Φ80Δ*lacM*15 Δ*lacX*74 *hsdR*($r_K^-$ $m_K^+$) | Invitrogen |
| Rosetta 2 (DE3) | $F^-$ *ompT hsdS*$_B$($r_B^-$ $m_B^-$) *gal dcm* (DE3) pRARE2 | Novagen/Merck |

* Δ10 mutations are described in detail in strain SA5666.

**Table S4.** Plasmids used in this work.

| Plasmid | Description | Reference |
|---|---|---|
| pBJ114 | *galK,* $Km^r$ | [35] |
| pSW105 | *attP,* $P_{pilA}$ $Km^r$ | [36] |
| pSWU30 | *attP,* $Tet^r$ | [37] |
| pMR3691 | *MXAN_18-19, vanR* $P_{van}$, $Tet^r$ | [38] |
| pMAL-c6t | Expression vector, $P_{tac}$, $His_6$-MalE, $Amp^r$ | NEB |
| pET28(+) | Expression vector, $P_{T7}$ $His_6$, $Km^r$ | Novagen |
| pMAT162 | pBJ114, in-frame deletion construct for *pilA,* $Km^r$ | [22] |
| pDK25 | pBJ114, in-frame deletion construct for *gltB,* $Km^r$ | [8] |
| pAP19 | pBJ114, in-frame deletion construct for *frzE,* $Km^r$ | [31] |
| pMP072 | pBJ114, in-frame deletion construct for *dmxA,* $Km^r$ | This work |
| pTP140 | pSWU30, $P_{nat}$ *dmxA,* $Tet^r$ | [28] |
| pMP092 | pSWU30, $P_{nat}$ *dmxA-mVenus,* $Tet^r$ | This work |
| pMP093 | pBJ114, *dmxA* replacement by *dmxA-mVenus*, $Km^r$ | This work |
| pMP164 | pBJ114, for native site integration of *dmxA-mVenus,* $Km^r$ | This work |
| pMP165 | pBJ114, for native site integration of *dmxA*$^{E626A}$*-mVenus,* $Km^r$ | This work |
| pMP095 | pBJ114, *dmxA* replacement by *dmxA*$^{R615A}$*-mVenus,* $Km^r$ | This work |
| pKH02 | pBJ114, in-frame deletion of *mVenus* in the *dmxA*$^{R615A}$*-mVenus* strain, $Km^r$ | This work |
| pMP175 | pBJ114, *dmxA* or *dmxA-mVenus* replacement by *dmxA*$^{\Delta GAF\times2\ (\Delta 223\text{-}532)}$, $Km^r$ | This work |
| pMP179 | pBJ114, *dmxA* or *dmxA-mVenus* replacement by *dmxA*$^{\Delta TMD\ (\Delta 3\text{-}222)}$, $Km^r$ | This work |
| pMP182 | pBJ114, for native site integration of *dmxA*$^{TMD\ (1\text{-}167)}$*-mVenus,* $Km^r$ | This work |
| pMH113 | pMAL-c6t, *malE-dmxA*$^{WT(223\text{-}722)}$, $Amp^r$ | This work |
| pTP139 | pET28a, $His_6$-*dmxA*$^{223\text{-}722,\ E626A}$, $Km^r$ | This work |
| pMH117 | pMAL-c6t, *malE-dmxA*$^{E626A\ (223\text{-}722,\ E626A)}$, $Amp^r$ | This work |
| pMP082 | pET28a, $His_6$-*dmxA*$^{223\text{-}722,\ R615A}$, $Km^r$ | This work |
| pMH116 | pMAL-c6t, *malE-dmxA*$^{R615A\ (223\text{-}722,\ R615A)}$, $Amp^r$ | This work |
| pMH114 | pMAL-c6t, *malE-dmxA*$^{GAF\times2\ (223\text{-}547)}$, $Amp^r$ | This work |
| pMH115 | pMAL-c6t, *malE-dmxA*$^{GGDEF\ (546\text{-}722)}$, $Amp^r$ | This work |
| pMAT74 | pBJ114, in-frame deletion construct for *ftsZ,* $Km^r$ | This work |
| pMAT86 | pMR3691, *ftsZ*, $Tet^r$ | This work |
| pMH52 | pEX-k168, *miniTurbo-FLAG*, $Km^r$ | This work |

| | | |
|---|---|---|
| pMP172 | pSW105, *dmxA-miniTurbo-FLAG* , Km$^r$ | This work |
| pMH98 | pBJ114, *ftsK* replacement by *ftsK-miniTurbo-FLAG*, Km$^r$ | This work |
| pMH110 | pSW105, *ftsK-miniTurbo-FLAG*, Km$^r$ | This work |
| pBBR15.2-2H12.D11opt-scarRef | pBBR15.2, $P_{tet\text{-}const}$ *cdGreen2 mScarlet-I*, Gm$^r$ | [30] |
| pMH123 | pSW105, *cdGreen2 mScarlet-I*, Km$^r$ | This work |
| pAP37 | pBJ114, *pilQ* replacement by *pilQ-sfGFP,* Km$^r$ | [31] |
| pMAT336 | pBJ114, *pilM* replacement by *mCherry-pilM,* Km$^r$ | [2] |
| pMEM33 | pBJ114, *pilT* replacement by *mCherry-pilT,* Km$^r$ | This work |
| pMEM23 | pBJ114, *pilB* replacement by *pilB-mCherry,* Km$^r$ | This work |
| pLC32 | pBJ114, *romR* replacement by *romR-mCherry,* Km$^r$ | [22] |
| pLC58 | pBJ114, *mglB* replacement by *mglB-mVenus,* Km$^r$ | [23] |
| pLC20 | pBJ114, *mglA* replacement by *mglA-mVenus,* Km$^r$ | [22] |
| pAP35 | pBJ114, *sgmX* replacement by *sgmX-mVenus,* Km$^r$ | [31] |
| pDJS01 | pBJ114, in-frame deletion construct for *dmxB* (*MXAN_3735*), Km$^r$ | [28] |
| pDJS02 | pBJ114, in-frame deletion construct for *MXAN_5366,* Km$^r$ | [28] |
| pDJS03 | pBJ114, in-frame deletion construct for *MXAN_7362,* Km$^r$ | [28] |
| pIH01 | pBJ114, in-frame deletion construct for *gacA* (*MXAN_4463*), Km$^r$ | [28] |
| pIH03 | pBJ114, in-frame deletion construct for *MXAN_5791,* Km$^r$ | [28] |
| pNGS010 | pBJ114, in-frame deletion construct for *MXAN_2997,* Km$^r$ | [28] |
| pTP120 | pBJ114, in-frame deletion construct for *MXAN_1525,* Km$^r$ | [28] |
| pTP121 | pBJ114, in-frame deletion construct for *MXAN_5199,* Km$^r$ | [28] |
| pTP125 | pBJ114, in-frame deletion construct for *gacB* (*MXAN_2643*), Km$^r$ | [28] |
| pTP127 | pBJ114, in-frame deletion construct for *MXAN_4029,* Km$^r$ | [28] |

**Table S5.** Oligonucleotides used in this work[1]

| Primer name | Sequence 5'-3' | | Brief description |
|---|---|---|---|
| 3702_map_fwd2 | TGACCACCTGGTCTGTACGC | Operon mapping Fragment A | Primers used for operon mapping. |
| 3703_map_rev2 | GGGGGCCATAGTCCTTCTGG | | |
| 3704_map_rev2 | TCATTGCGCAGGGCCTCATT | Operon mapping Fragment B | |
| 3703_map_fwd2 | CGCGAACCAGGCGATGATTG | | |
| 3704_map_fwd2 | AGGCCCTGCGCAATGAGATT | Operon mapping Fragment C | |
| 3705_map_rev2 | AATGGCCACCACGACGAAGG | | |
| 3705_map_fwd2 | GCCGCTGAGGATCACCATGT | Operon mapping Fragment D | |
| 3706_map_rev2 | CTCTGGGGCATCGGCCTG | | |
| ftsZ-up EcoRI | GCGCGAATTCGGTGGACACGGATGGCGACG | | For deletion of *ftsZ* |
| ftsZ-overlapping reverse | CGTCTGGCCCTTGTTCTGATCGAACTGGTC | | |
| ftsZ-overlapping forward | CAGAACAAGGGCCAGACGGAACTGCCGTAA | | |
| ftsZ-down HindIII | GCGCAAGCTTTTGCGCAGCTGCCAGCCGATG | | |
| ftsZ-start NdeI | GGAATTCCATATGGACCAGTTCGATCAGAAC | | For expression of *ftsZ* under the $P_{van}$ promoter |
| ftsZ-stop KpnI | GCGCGGTACCTTACGGCAGTTCCGTCTGGC | | |
| sfGFP_fwd_NdeI | GCGCCATATGAGCAAAGGAGAAGAACT | | For expression of *sfGFP-miniTurbo-FLAG* under the $P_{van}$ promoter |
| sfGFP_rev_OV | CGCCCCCGCCTTTGTAGAGCTCATCCATGC | | |
| TurboID_fwd_OV | GCTCTACAAAGGCGGGGGCGGGAGCATGAT | | |
| TurboID_rev_EcoRI | GCGCGAATTCTCACTTGTCGTCGTCGTC | | |
| FtsK_A_BamHI | GCGCGGATCCACGAGGACGACATGCTCGACGC | | For replacement of *ftsK* with *ftsK-miniTurbo-FLAG* |
| FtsK_B_TID_OV | CGCCCCCGCCCATGGCCCCGGCGCCGGGCA | | |
| TID_fwd_FtsK_OV | CGGGGCCATGGGCGGGGGCGGGAGCATGAT | | |
| TID_rev_FtsK_OV | CACGTGGGCCTCACTTGTCGTCGTCGTCCT | | |
| FtsK_C_TID_OV | CGACAAGTGAGGCCCACGTGGCCGCCGTCT | | |
| FtsK_D_HindIII | GCGCAAGCTTCACCTTGAGGGCCACGAAGCCGG | | |
| FtsK_fwd_XbaI_2 | GCGCTCTAGACAGCCGAAATGTAGGTTCCCCGTCCTGAGC | | For expression of *ftsK-miniTurbo-FLAG* from the *attB* site and $P_{pilA}$ |
| FtsK_B_TID_OV 2 | CGCCCCCGCCCATGGCCCCGGCGCCGGGCATGTCG | | |
| TID_fwd_FtsK_OV | CGGGGCCATGGGCGGGGGCGGGAGCATGAT | | |
| Flag_rev HindIII | GCGCAAGCTTTCACTTGTCGTCGTCGTC | | |
| MalE-fwd_NotI | GCGCGCGGCCGCGAGATCGAAGGCGCCGTGC | | For expression and purification of MalE-tagged DmxA variants |
| DmxA_Rev_HindIII | GCGCAAGCTTTCAGGACGCGTTCGCCGCCT | | |
| DmxA_GAF2_rev_HindIII | CTGAGGATCCTCACGTGGTGGCCATGCGCTCCA | | |
| DmxA_GGDEFonly_Fwd_NotI | CTGAGGATCCTCACGTGGTGGCCATGCGCTCCA | | |
| 3705 GAF1 forw NdeI | ATCGCATATGGAGATCGAAGGCGCCGTGC | | For expression and purification of His6-tagged DmxA protein variants |
| 3705 E626A (-) | GACGAACTCCGCGCCGCCGTA | | |
| 3705 E626A (+) | TACGGCGGCGCGGAGTTCGTC | | |
| 3705 R615A (-) | GTCCGTATCGGCCGCCATCGTC | | |
| 3705 R615A (+) | GACGATGGCGGCCGATACGGAC | | |

| | | |
|---|---|---|
| 3705 rev BamHI | ATCGGGATCCTCAGGACGCGTTCGCCGC | |
| cdG-Sensor_fwd_XbaI | ATGCTCTAGAATGAATTCGGAACCGCCGCC | For expression of cdGreen and mScarlet-I from the *attB* site and the $P_{pilA}$ |
| cdG-Sensor_rev_HindIII | ATTCAAGCTTTCACTTGTACAGTTCATCCATACCACCG | |
| 3705_A | ATCGGGTACCAAGGTGACGTCGCACAAGAT | For deletion of *dmxA* |
| 3705_B | CTTCAGCTGAGACGGAAACTCGGGAAG | |
| 3705_C | TTTCCGTCTCAGCTGAAGGCGGCGAAC | |
| 3705_D | ATCGTCTAGACATCACCGGGTGGACCGTCA | |
| 3704 prmt forw +XbaI | ATCGTCTAGATTCCTCAACGCGCTGGCGCTG | For $P_{nat}$ *dmxA-mVenus* |
| 3705_rev no stop 1 | GCCGCCGCCGGACGCGTTCGCCGCCTTCAG | |
| 3705_mVenus fw | AACGCGTCCGGCGGCGGCGGCTCCATGGTGAGCAAGGG | |
| mVenus_Kpn rev | CGCGCCGGGTACCTTACTTGTACAGCTCGTCCA | |
| 3705_native forw | ATATGGTACCTCCGCTACGGGGCGCCGCTG | For replacement of *dmxA* with *dmxA-mVenus* |
| 3705_native middle rev | CCCGGCTGCATCAAGGACTTACTTGTACAGCTCGTC | |
| 3705_native middle fw | GACGAGCTGTACAAGTAAGTCCTTGATGCAGCCGGG | |
| 3705_native rev | CGCGTCTAGACATCCCCGAGTCGGCCATCG | |
| 3705_PpilA forw | CGCGTCTAGAATGACGTCGCTTCCCGAGTT | For $P_{pilA}$ *dmxA-miniTurbo-FLAG* |
| 3705_miniTurboID fw | AACGCGTCCGGCGGCGGCGGCTCCATGATCCCGCTCCTGA | |
| 3705_B (GAFx1) | CGCGCGCAGCTTCACCGAGGCCACCAG | For in-frame deletion of the GAF domains of *dmxA* |
| 3705_C (GAFx2) | TCGGTGAAGCTGCGCGCGCAGCTCTAC | |
| 3705_D (GAFx2) | ATCGGGTACCGGACGCGTTCGCCGCCTTCA | |
| 3705_B3 (TMD) | TTCGATCTCCGTCATGGGGACTCCAGG | For native site integration of *dmxA*$^{\Delta TMD}$ *-mVenus* |
| 3705_C3 (TMD) | CCCATGACGGAGATCGAAGGCGCCGTG | |
| 3705_D (TMD)2 | ATCGTCTAGACAGCGGCGCCCCGTAGCGGA | |
| 3705_TMH rev | GGAGCCGCCGCCGCCCAGCACCAGGTGGTAGAGGC | For native site integration of *dmxA*$^{TMD\ (1\text{-}167)}$*-mVenus* |
| 3705_mVenus fw2 | CTGGTGCTGGGCGGCGGCGGCTCCATGGTGAGCAAGGG | |
| 3705_del_mVenus_(-) | AAGGACTCAGGACGCGTTCGCCGCCTTCAG | For deletion of *mVenus* |
| 3075_del_mVenus_(+) | AACGCGTCCTGAGTCCTTGATGCAGCCGGG | |
| pilB-mCA-EcoRI | GCGCGAATTCGATCACCTCGCGTTTGAAG | For replacement of *pilB* with *pilB-mCherry* |
| pilB-mCB-overlay | GAGGAAGGTTGACTACTTGTACAGCTCGTCCATG | |
| pilB-mCC+ overlay | GACGAGCTGTACAAGTAGTCAACCTTCCTCCCAC | |
| pilB-D-HindIII | GCGCAAGCTTGGTCTGCATGCCGAACTTCG | |
| PilT fw EcoRI | CGCGAATTCGCTCCTTCCTCC | For replacement of *pilT* with *mCherry-pilT* |
| PilT rev HindIII | GCGCAAGCTTCTAACGACCAC | |

[1] Underlined sequences indicate restriction sites.

**Table S6.** Fully sequenced myxobacterial genomes used for the 16S rRNA tree

| Species and strain name |
|---|
| *Anaeromyxobacter dehalogenans* 2CP-C |
| *Anaeromyxobacter* sp. Fw109-5 |
| *Anaeromyxobacter* sp. K |
| *Anaeromyxobacter oryzae* Red232 |
| *Anaeromyxobacter paludicola* Red630 |
| *Archangium gephyra* DSM 2261 |
| *Archangium violaceum* SDU34 |
| *Chondromyces crocatus* Cm c5 |
| *Corallococcus macrosporus* DSM 14697 |
| *Corallococcus coralloides* DSM 2259 |
| *Cystobacter fuscus* DSM 52655 |
| *Haliangium ochraceum* DSM 14365 |
| *Labilithrix luteola* DSM 27648 |
| *Melittangium boletus* DSM 14713 |
| *Minicystis rosea* DSM 24000 |
| *Myxococcus fulvus* 124B02 |
| *Myxococcus hansupus* (*Myxococcus* sp. mixupus) |
| *Myxococcus stipitatus* DSM 14675 |
| *Myxococcus xanthus* DK1622 |
| *Nannocystis* sp. fl3 |
| *Polyangium aurulentum* SDU3-1 |
| *Sandaracinus amylolyticus* DSM 53668 |
| *Sorangium cellulosum* So ce 56 |
| *Stigmatella aurantiaca* DW4/3-1 |
| *Vulgatibacter incomptus* DSM 27710 |
